# A Hybrid Residual–Swin Transformer Design with Attention for Prostate Cancer Segmentation

**DOI:** 10.64898/2026.09.14.751449

**Authors:** Rahul Singh, Sheifali Gupta, Sapna Juneja, Deepali Gupta, Sunil Maggu, Mingqiang Wang, Saurav Mallik

**Author notes:** **Corresponding Authors:** Sapna Juneja; Saurav Mallik (,).

## Abstract

Prostate cancer, a leading cause of cancer-related deaths among men globally, necessitates the development of precise diagnostic and treatment strategies. Accurate segmentation of prostate cancer in medical imaging, particularly in MRI scans, is crucial for early diagnosis and clinical decisions. Traditional manual segmentation techniques, while efficient, are labor-intensive and necessitate significant expertise, resulting in an increasing demand for automated alternatives. The RSAUNet is a novel deep learning architecture designed to improve prostate cancer segmentation. It incorporates essential components, including Residual Blocks, Swin Transformer Blocks, and Attention Mechanisms within a U-Net architecture. This markedly enhances the model’s capacity to discern complex anatomical features and accurately segment malignant tissues. Using sophisticated deep Learning methods, RSAUNet addresses the complexities of prostate imaging, delivering reliable, consistent segmentation results. The model was evaluated against various cutting-edge techniques on extensive multi-parametric MRI datasets, attaining an impressive Dice coefficient (DC) of 0.998 and a Jaccard Index (IoU) of 0.965. These findings highlight the innovative characteristics of RSAUNet and its capacity to transform prostate cancer diagnosis and treatment strategies. The proposed model surpasses current methods and shows potential for practical clinical applications, providing an efficient, precise, and scalable solution for automated prostate cancer segmentation and fostering optimism for the future of medical imaging and diagnosis.

## Introduction

Prostate cancer remains one of the most frequently diagnosed malignancies among men worldwide and continues to pose a major public health challenge due to its high prevalence and associated mortality risk [1,2]. Current estimates suggest that approximately one out of every eight men will develop prostate cancer during their lifetime [3]. Owing to its widespread occurrence, it is recognized as the second most commonly diagnosed cancer in men after non-melanoma skin cancer. In the United States, prostate cancer accounts for nearly 27% of all newly reported cancer cases among males [4]. Furthermore, annual statistics indicate that more than 200,000 new cases are diagnosed each year, highlighting the substantial burden of this disease on healthcare systems and affected populations [5]. With the average diagnosis age of approximately 65 years, this disease almost purely targets elderly males [3]. Prostate cancer is highly prevalent, which highlights the need for more in-depth studies and advanced diagnostic and treatment technologies. It is a disease that originates in the prostate, a small gland within the male reproductive structure in front of the rectum and below the bladder [6]. The principal role of the prostate is to secrete seminal fluid-which feeds and carries sperm [7, 8]. Generally, cancers of the prostate start as small, localized tumors called adenocarcinomas, and it is surmised that they originate in the epithelial cells of the prostate. These cancers may affect distant organs, such as lymph nodes and bones, after a long period of time [9]. Most prostate cancers take a slow course. This has made it possible to use a range of treatments that come from passive follow-up as active surveillance to more aggressive hormone therapy, radiation treatment, and surgery. Several known risk factors increase the possibility of prostate cancer. The biggest risk factor for this disease is age, and the incidence sharply rises after 50 years [10]. Men with a family history of prostate cancer are more than twice as likely to develop the condition than individuals from the general population, suggesting that genetics also play an important role [11]. Besides, some genetic mutations, for example, BRCA1 and BRCA2, have been associated with an increased risk of prostate cancer, particularly aggressive types of prostate cancer [6]. Ethnically, African American men are also diagnosed with a higher rate of prostate cancer than other races and also have the highest death rate with the most probable diagnosis of advanced prostate cancer [12]. These disparities have multiple causes ranging from genetics to environmental factors, and socioeconomic issues. The other lifestyle factors found to be associated with increased risk of prostate cancer are diet and physical activity. There is an association between diets low in fruits and vegetables and high in red meat or fat dairy products and prostate cancer. To the contrary, frequent exercise and a diet rich in fish, cruciferous vegetables and tomatoes can help to decrease risk [13].

Prostate cancer can be treated as surgery, radiation therapy and hormone therapy [14]. In general, men with localized (confined) prostate cancer that is considered low risk may be advised to undergo active surveillance. This approach to management involves frequent surveillance of cancer using PSA tests, DREs and routine biopsy to confirm that the cancer is stable [15]. The main advantage of this technique is to prevent or postpone the possible complications of more aggressive therapies such as urinary and sexual impotence. Watchful waiting is the main treatment in older men or those with serious health problems, and focuses on symptom relief rather than eradication of the disease [3,9].

Medical imaging is an excellent diagnostic tool that can be used for staging and to develop treatment plans for prostate cancer. Decoding the data acquired from different medical images is difficult as the anatomy of prostate is generally intricate and the slight discrepancy between normal and cancerous tissue is very complex [16]. Segmentation of images is vital for the accurate and reliable diagnosis of prostate cancer by partitioning images into different regions of their images that correspond to anatomic structures and abnormalities. The classical techniques for image segmentation, among them thresholding algorithms and region-growing algorithms, are virtually applied in prostate imaging. These techniques, however, have definite challenges when applied, especially on variations in size, shape, and intensity. Advanced segmentation approaches have been developed due to these limitations, among them being the atlas-based and the deformable models. These methods use anatomical priors and deformable models to improve the accuracy of the segmentation. However, they require too much human input and knowledge.

To respond to some of the deficiencies in prostate cancer segmentation, an emerging model integrates advanced technologies, namely, RASUNet: Residual Attention Swin UNet. Many state-of-the-art technologies are implemented to enhance performance in segmenting structures involved in prostate cancer. More specifically, RASUNet aligns residual convolutional blocks, swin transformer blocks, and attention mechanisms with UNet architecture. Residual blocks enable the gradient flow and allow for the development of much deeper networks without suffering from the vanishing gradient, which is required for accurate feature extraction in complex anatomical regions such as the prostate. The Swin Transformer blocks use a global attention mechanism, which would help distinguish between cancerous and noncancerous tissues. The attention mechanisms boost the model’s concentration for accurate and context-aware segmentation. The contributions of the proposed model are clearly demonstrated by its advantages in the generalization ability across different medical datasets, and its high accuracy when applying these models for prostate cancer segmentation or other difficult segmentation problems. The contributions of this research work are as follows:

1. The model applies residual convolutional blocks in both the encoder and decoder, which improve feature extraction while alleviating the vanishing gradient problem. These residual connections help the model maintain better gradient flow through deeper layers, resulting in more robust learning, especially for complex tasks like segmentation.
2. The model incorporates Swin Transformer blocks within the encoder and bridge sections. These blocks utilize window-based self-attention mechanisms to capture both local and global dependencies. This allows the model to better focus on relevant spatial regions while maintaining context across larger areas, making it highly effective in distinguishing between similar-looking regions in medical images.
3. Attention mechanisms are applied in the decoder path, which enhance the model’s ability to focus on important regions of interest during upsampling. By weighting features based on their significance, the model can better capture critical structures in the image, leading to more accurate segmentation outputs.

The paper can be divided into Section 2, recent developments on prostate cancer segmentation, which is a deep learning-based application, mostly utilized in the medical imaging domain. Large-scale MRI data set for multi-parametric MRI images from Radboud University is described. This work explores a new architecture of RSAUNet by incorporating residual blocks and Swin Transformer blocks along with their attention mechanisms inside the UNet backbone. Section 4 describes the experimental work as the RSAUNet compares to the state-of-the-art methods, including quantitative and qualitative, such as the DC and Jaccard Index (IoU). Section 5 outlines significant findings, provides some paths for potential future research tracks, and suggests possible RSAUNet applications to the broader medical imaging community.

### Literature Review

Jiang et al. [21] applied images obtained from micro-ultrasound of 75 patients to segment the prostate. The architecture proposed employed multi-scale deep supervision with TransUNet and achieved a Dice coefficient of 0.939. Such research indicates that the accuracy in the process of segmentation of prostate regions can be very high and may greatly be instrumental in advancing the technology for images involved. Zhang et al. [22] presented using the PI-CAI dataset for developing an SFHGE-based, CrossTA, and HIFN-based segmentation-assisted prostate cancer grading model. The developed model achieved an 85.7% success rate, meaning efficient performance in prostate cancer grading and supporting procedure diagnosis. By using the Prostate MR Image Database, Prasad et al. [23] developed a GBVO U-Net++ model coupled with DCNN optimized using Gradient Bald Eagle Optimization (GBEO). The model achieved an accuracy of 0.916, a FNR of 0.104 and a FPR of 0.100, meaning improved accuracy on prostate cancer segmentation and detection, hence suggesting a new approach to improve prediction performance. Wang et al. [24] worked with 25 men with high-risk prostate cancer with the help of PET/CT and the tracer 18F-DCFPyL. They utilized semiautomatic segmentation, using both SUV% threshold and adaptive segmentation approaches. They demonstrated that threshold A40% was capable of providing a median relative difference of 17.6% for TV-Histo, and thus that was the best approach to ensure reliable segmentation in case of PET/CT imaging for prostate cancer. Toosi et al. [25] developed a self-supervised 3D segmentation approach by integrating DDPM, MA-MIP, and the OSEM Algorithm for automatic 3D segmentation from a dataset of 510 PSMA-PET/CT images. The model obtained an average value of the DC of 0.532 and an average value of the Jaccard index at 0.433, indicating potential improvements in PET imaging for the segmentation of prostate cancer. Kou et al. [26] applied their prostate cancer segmentation model on the Prostate158 and PROSTATEx2 datasets with ViT-base Model, and 95%HD. The results indicated that the DSC values were 83.2 ± 8.9 for Prostate158 and 81.0 ± 11.6 for PROSTATEx2, showing that the model is robust and versatile on different datasets thus ensures reliable segmentation in clinical settings. Jeong et al. [27] was developed to analyze CT images from 18 prostate cancer patients, who introduced a U-Net-based architecture (nnU-net) for radiation therapy in prostate cancer. The performance of the model was with DSC values of 0.94 for the bladder, 0.84 for the prostate and 0.83 for the rectum, it represented high accuracy and thus presented applicability in radiation therapy treatment planning. Jafari et al. [28] developed a CNN-based approach to the automatic whole-body [68Ga] Ga-PSMA PET images from 412 patients using a 3D CNN along with the framework of nnU-Net. The model reported accuracy at 83% for patient-level classification, 87%-94% for lesion-level recognition, and voxel-level segmentation with the DC between 65%-70%, which is pretty good, especially for large-scale clinical PET imaging. Kuanar et al. [29] proposed deep learning zonal segmentation using nnU-Net, leveraging the dataset of 1020 prostate MRIs, internal testing using 3461 test cases and external testing using 1460 test cases. This model used PSA-density as well as transition zone PSAD (TZ-PSAD), achieving Dice scores of 0.91 for TZ as well as 0.85 for PZ along with AUC scores of 0.75 and 0.76 respectively for TZ-PSAD on the internal and external test sets, respectively. This study showed improved segmentation and detection accuracy of clinically relevant prostate cancer. Li et al. [30] used a dataset of 133 patients, in which 71 patients had prostate cancer and 62 patients had benign tumors, for the task of prostate cancer segmentation using 3D-Mask RCNN. The method resulted in a DSC of 0.856 in the training set, along with 0.921 sensitivity and 0.961 specificity. The model scored a DSC of 0.849 on the testing set while producing sensitivity and specificity measures of 0.911 and 0.931, respectively. Alzate-Grisales et al. [31] used SAM-UNETR, which was the SAM transformer encoder applied to a residual-convolution UNETR decoder. It enabled transfer learning onto the Prostate158 and PI-CAI Challenge data. The model resulted in a Dice score (DS) of 0.624 and an IoU score of 0.491, showing the complexity involved with large prostate cancer segmentation using transformers. Pellicer-Valero et al. [32] proposed a 3D Retina U-Net model. A mean DSC was reported as 0.945, meaning the model is quite accurate in the segmentation of prostate cancer and diagnostic support. Duran et al. [33] developed a ProstAttention-Net, an attention mechanism incorporated network based on the dataset given by 219 MRI scans in PROSTATEx-2. The model attained a DS of 0.875 with a strong segmentation framework improved by attention mechanisms that could support accurate prostate segmentation and cancer diagnosis. Zhang et al. [34] presented the cross-modal framework using self-attention distillation and spatial correlated feature fusion (SCFF) techniques on a dataset consisting of 358 MRI images. Their method resulted in a DSC value of 65.7 ± 1.2, which promised but still required further development to improve the segmentation. Chahal et al. [35] proposed a UNet-based Xception model using the PROMISE12 dataset. The model used local residual connections in the decoder. With a Dice coefficient of 97.50%, it presented an excellent model for identifying regions of prostate from MRI images. Kiljunen et al. [36] presented the AST model to the anatomy segmentation of 30 patients. In their methodology, they attained a Dice similarity coefficient of 0.94, becoming an effective tool in automated anatomy segmentation in radiation therapy planning with prostate cancer. Zhang et al. [37] designed a bi-attention adversarial network by utilizing MR T2-weighted images in a cohort of 120 patients. The model achieved 86.4% accuracy, 93.2% sensitivity, 75.7 Jaccard Index, and 85.9 Dice coefficient, providing a strong foundation for prostate cancer segmentation by adversarial learning. Hossain et al. proposed VGG19RSeg, a residual semantic segmentation model for prostate MR images. The DSC reached an acceptable value of 94.57% in using VGG19RSeg thereby surpassing the accuracy of similar methods and boosting the precision of prostate cancer segmentation.

**Table 1:** Literature Review.

| Ref/Year | Dataset Used | Techniques | Results |
| --- | --- | --- | --- |
| [21]/2024 | Micro-ultrasound images of 75 patients | Multi-scale deep supervision, TransUNet architecture | Dice coefficient: - 0.939 |
| [22]/2024] | PI-CAI Dataset | Shared Feature Hybrid Gating Experts (SFHGE), Cross-Task Attention (CrossTA), Heterogeneous Information Fusion Network (HIFN) | Accuracy: - 85.7% |
| [23]/2024 | Prostate MR Image Database | GBVO-based U-Net++ with DCNN, Gradient Bald Eagle Optimization (GBEO) | Accuracy: - 0.916<br>False Negative Ratio: - 0.104<br>False Positive Ratio: - 0.100<br>Negative Predictive Value: - 0.903 |
| [24]/2024 | PET/CT imaging with 18F-DCFPyL | Semiautomatic segmentation using SUV% thresholds (30%-70%) and adaptive segmentation (A30%-A70%) | Adaptive segmentation with A40% threshold had a median relative difference of 17.6% from TV-Histo and was most accurate compared to other methods. |
| [25]/2024 | 510 PSMA-PET/CT images | Diffusion models (DDPM), Multi-angle Maximum Intensity Projections (MA-MIP), OSEM Algorithm | Dice Coefficient: - 0.532<br>Jaccard: - 0.433 |
| [26]/2024 | Prostate158, PROSTATEx2 | Coarse segmentation U-Net, Rejection Network, SAM, ViT-base Model, 95%HD | **Prostate158**<br>Dice similarity coefficient (DSC): - $83.2 \pm 8.9$<br>True Positive Rate: - 91.7<br>Positive Predictive Value: - 91.8<br>F1- Score: - 91.7<br>**PROSTATEx2**<br>DSC: - $81.0 \pm 11.6$<br>True Positive Rate: - 93.5<br>Positive Predictive Value: - 88.3<br>F1- Score: - 90.8 |
| [27]/2024 | CT images of 18 prostate cancer patients | U-Net-based architecture (nnU-net) | Bladder DSC: - 0.94<br>Prostate DSC: - 0.84<br>Rectum DSC: - 0.83 |
| [28]/2024 | 412 patients from three centers ([68Ga] Ga-PSMA PET/CT scans) | nnU-Net framework, 3D convolutional neural network (CNN), patch-based segmentation | Voxel-level segmentation<br>DSC: - 65%-70%<br>PPV: - 72-79%<br>IoU: - 53-58%<br>Sensitivity: - 62-73% |
| [29]/2024 | 1020 prostate MRIs for model development, 3461 additional internal test cases, and 1460 external test cases (PI-CAI challenge) | Deep learning zonal segmentation using nnU-Net, PSA-density (PSAD), transition zone PSAD (TZ-PSAD) | Dice scores: 0.91 (TZ), 0.85 (PZ);<br>AUC: 0.75 (TZ-PSAD), 0.73 (PSAD) on internal test set, 0.76 (TZ-PSAD), 0.74 (PSAD) on external test set |
| [30]/2023 | 133 patients (71 with prostate cancer, 62 benign tumors), T2WI MRI | 3D-Mask RCNN | Training set<br>DSC: 0.856,<br>Sensitivity: 0.921,<br>Specificity: 0.961.<br>Testing set<br>DSC: - 0.849<br>Sensitivity: - 0.911<br>Specificity: - 0.931 |
| [31]/2023 | Prostate158, PI-CAI Challenge datasets | SAM-UNETR (combines SAM transformer encoder with residual-convolution UNETR decoder, transfer learning) | Dice Score: - 0.624<br>IoU: - 0.491 |
| [32]/2022 | ProstateX, Valencian Oncology Institute Foundation | 3D Retina U-Net | Mean DSC: -0.945 |
| [33]/2022 | 219 MRI (PROSTATEx-2) | ProstAttention-Net with an attention mechanism | Dice score: - 0.875 |
| [34]/2021 | 358 MRI images | SCFF | DSC: - $65.7 \pm 1.2$<br>Sensitivity: - $77.8 \pm 2.5$<br>Precision: - $60.7 \pm 0.4$ |
| [35]/2021 | PROMISE12 Dataset | UNet with Xception in Encoder, Local Residual Connections in Decoder | Dice coefficient: - 97.50% |
| [36]/2020 | 30 Prostate Cancer Patients | DL-based AST model | DSC: - 0.94 |
| [37]/2019 | MR T2-weighted images from 120 patients from Shanghai Tenth People Hospital | Bi-attention adversarial network (GAN with U-Net as the backbone) | Accuracy: - 86.4<br>Sensitivity: - 93.2<br>Jaccard Index (IoU): - 75.7<br>Dice Coefficient: - 85.9 |
| [38]/2018 | ----- | VGG19RSeg | DSC: - 94.57% |

### Proposed Methodology

The proposed methodology for prostate cancer segmentation integrates three key components - Residual Blocks, Swin Transformer Blocks, and Attention Mechanism Blocks - to improve segmentation accuracy, as shown in Figure 1. To address vanishing gradients and ensure the information passes through, the Residual Block is created with shortcut connections, which add the input to the output of the block, so the model can learn low-level features well. The Swin Transformer Block adds global context awareness by incorporating window-based multi-head self-attention (MHSA) and normalization as well as multi-layer perceptron (MLP) layers, with the addition of residual connections to preserve important information that enhances the ability to capture complex patterns. Lastly, the Attention Mechanism Block highlights the relevant regions of the image through convolutional layers, ReLU and sigmoid activations to assign weight to features that are relevant to the segmentation task. Together, these components enable RSAUNet to achieve high performance in segmenting prostate zones, which is crucial for diagnosing and treating prostate cancer.

**Figure 1:**
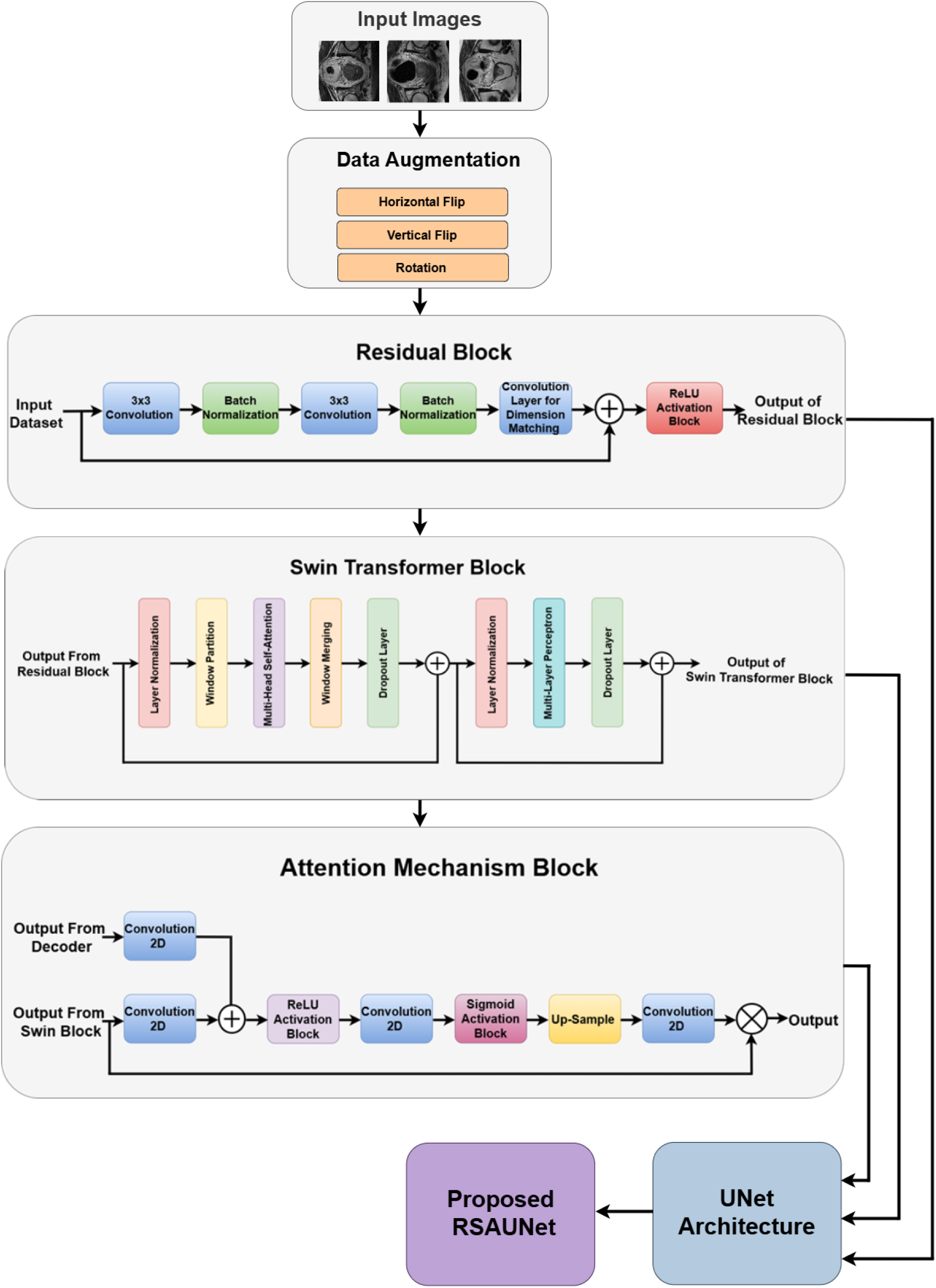
Proposed Workflow.

### Input Dataset

For experimentation purposes, a publicly available dataset available at Kaggle [39] has been used. High-resolution MRI scans of the transverse T2-weighted images (T2WIs) from a 0.6 x 0.6 x 4 mm resolution and apparent diffusion coefficient (ADC) maps from a 2 x 2 x 4 mm resolution have been acquired, enabling the detailed anatomical characterization of the prostate and tissue differentiation. An extremely accurate manual segmentation of the prostate was carried out to generate pixel-wise ground truth masks for training and evaluation of deep learning models. Some typical examples from the dataset are shown in Figure 2; each row depicts an input MRI image and the corresponding manually annotated segmentation mask. The examples, therefore, demonstrate the anatomical variation of presentation and difficulty in differentiation from adjacent tissues, thus making it suitable for preparing a robust and accurate segmentation model. Using input images and true masks is helpful to visually verify the model’s effectiveness in correct outline delineation of the prostate for applications in diagnosing prostate cancer and planning for treatment.

**Figure 2:**
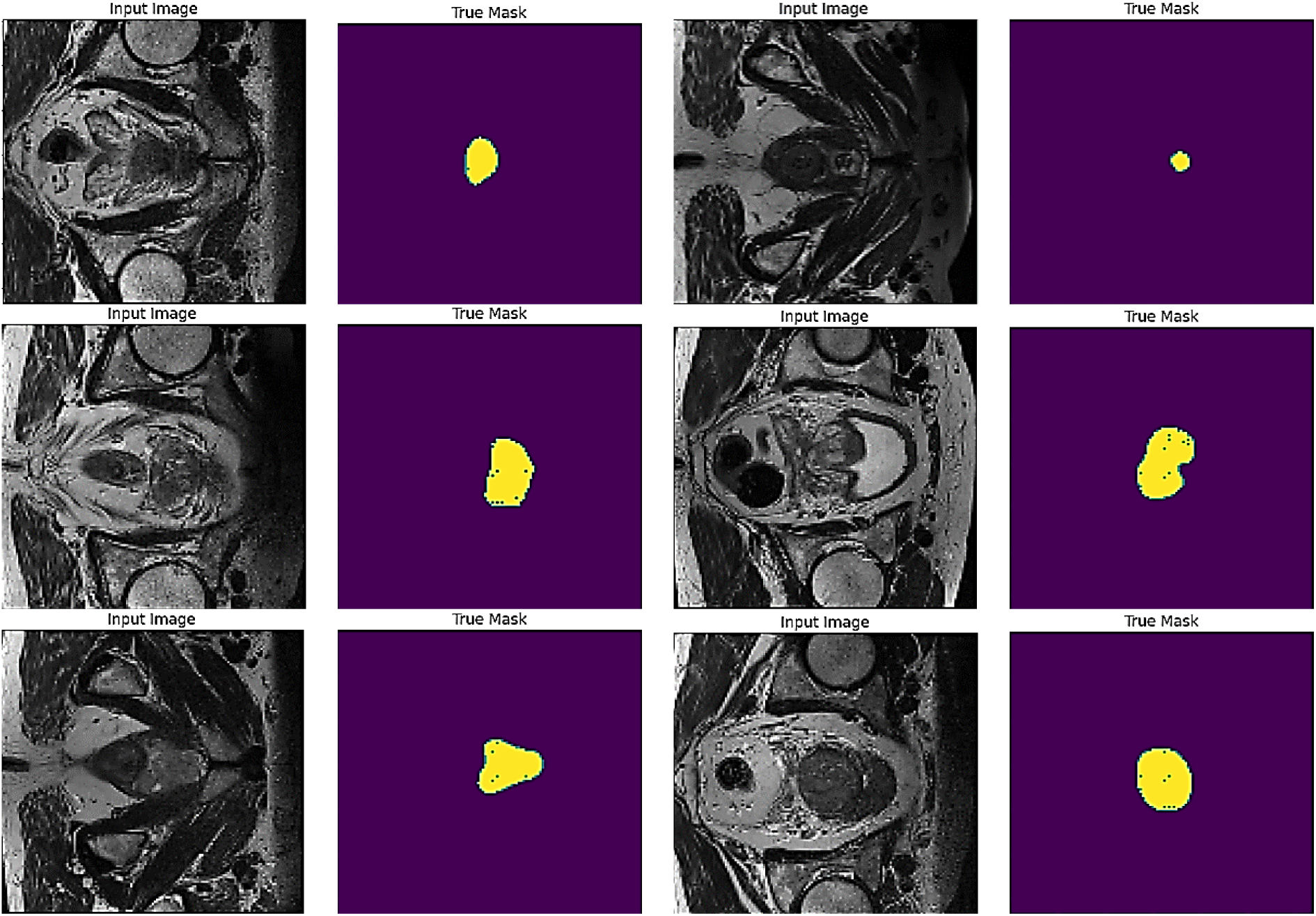
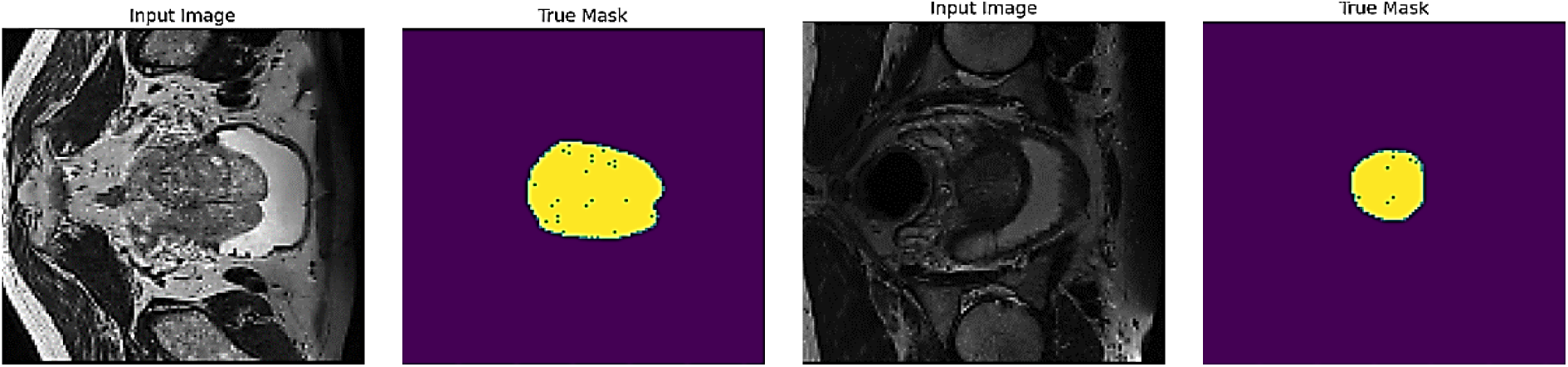
Input Dataset.

### Data Augmentation

Figure 3 displays augmentation performed on the prostate segmentation dataset. The augmentations used were horizontal flip, vertical flip, and rotation, as shown in Figures 3(b), 3(c), and 3(d). Usually, flips and rotations are deep learning standard augmentations for the generalization and robustness of models. All these transformations would be applied to the original images, expanding the variability of the images and enabling the model to learn from various views and orientations of the prostate (see Figure 3(a) for original images). This would reduce overfitting and increase the odds of the model being able to do a good job in segmenting new prostate data. The augmentation helps to simulate different clinical situations and ensure the model’s performance is consistent under different image settings.

**Figure 3:**
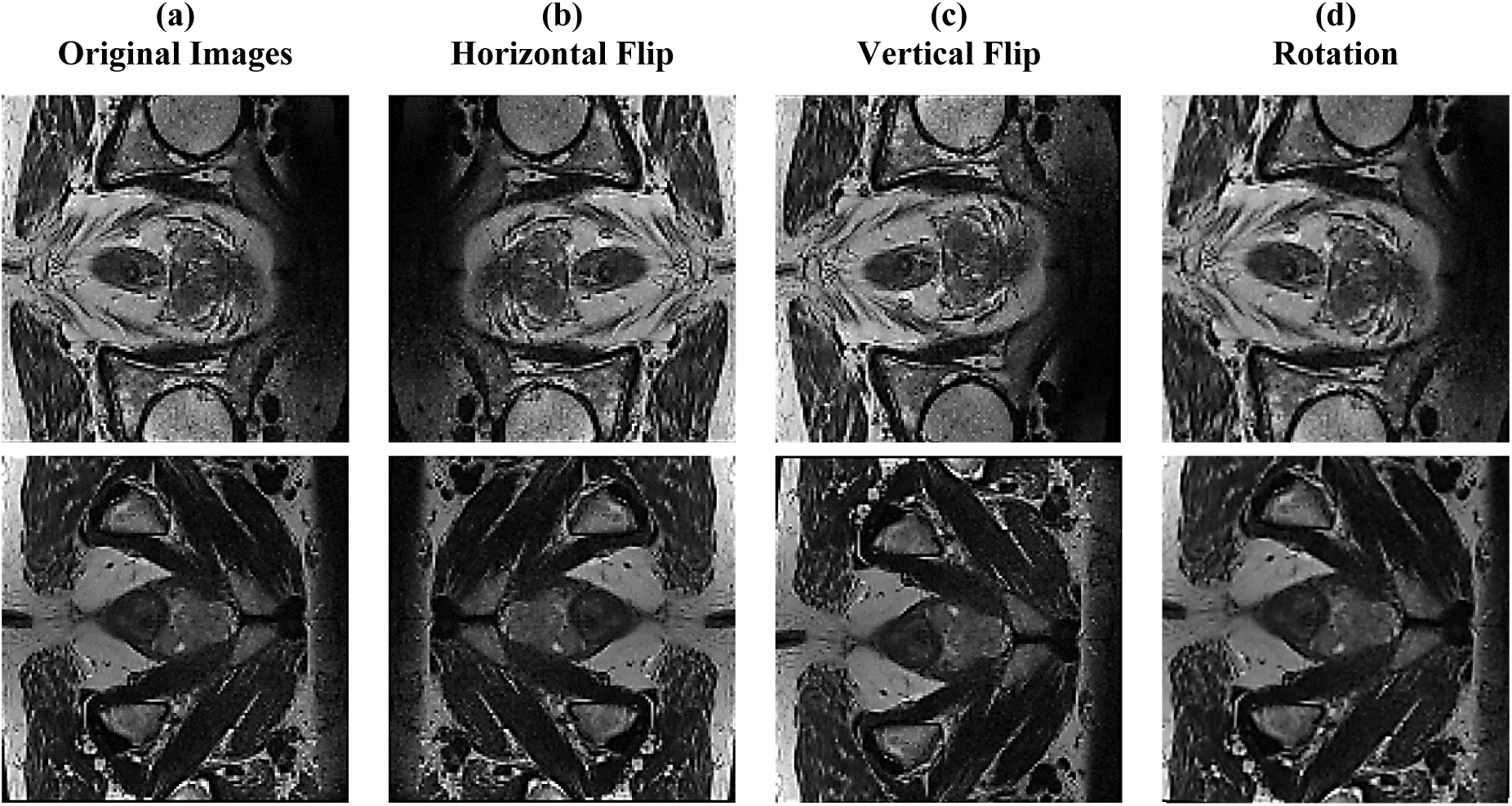
Data Augmentation: (a) Original Images, (b) Horizontal Flip, (c) Vertical Flip, (d) Rotation.

### Residual Block

This is an essential part of deep neural networks, especially regarding the vanishing gradient problem. The block has an initial input dataset, passed on to two successive layers, each of which is a 3x3 convolution and followed by Batch Normalization to improve stability and speed in training. Both are intended to excerpt intricate structures from the input data. A traditional convolution layer, colored blue, ensures its output dimensions match the input dimension for the addition. The core of a residual learning framework is the shortcut connection, represented by the direct line bypassing the convolution and normalization layers that enable the addition of the output of the original input directly with the processed output. This addition is the core of the residual learning concept as it allows learning identity mappings that avoid the vanishing gradient problem. This sum now goes into a ReLU activation function that brings the non-linearity to the output; hence, the Residual Block output is produced. Therefore, this helps the network train deeper layers, effectively improving the model’s performance. Figure 4 shows the Residual Block.

**Figure 4:**
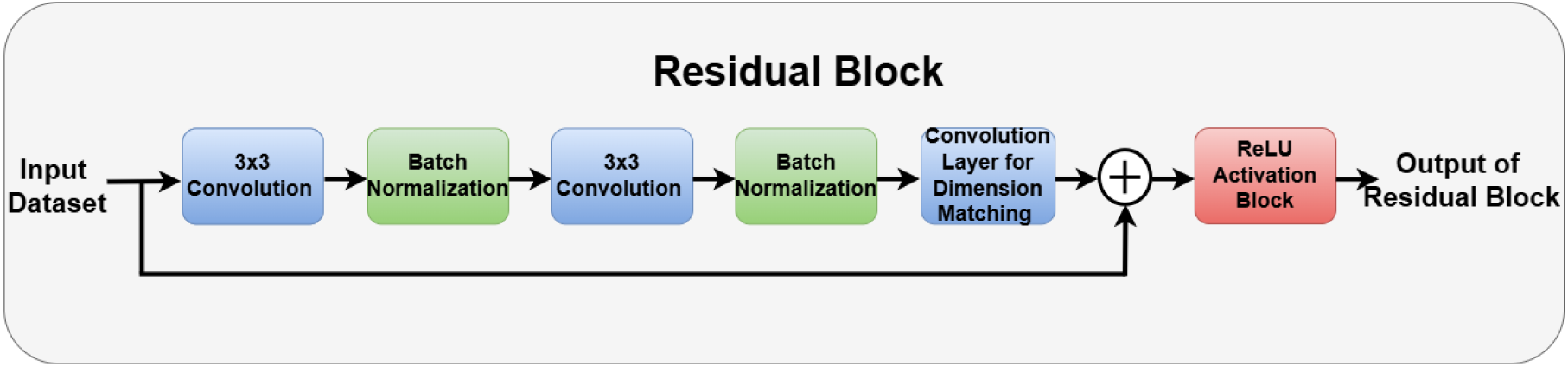
Residual Block.

### Equations and Operations

The mathematical representation of a Residual Block is given in equation 1, Where: *X* is the input to the Residual Block, *Conv* denotes a convolution operation, *BN* denotes batch normalization and *ReLU* denotes the ReLU activation function.

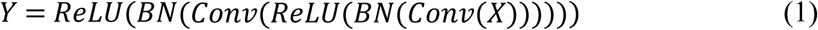

The Residual connection adds the input X back to the output of the convolutions. The equation of residual connection is given in equation 2:

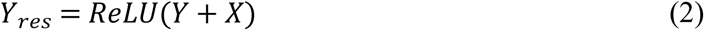

Here, *Y* + *X* represents the shortcut connection that adds the input directly to the output of the convolution layers. This addition helps learn identity functions, making the model’s learning easier even with very deep networks.

### Swin Transformer block

It is designed to improve the self-attention mechanism by implementing a shifted window mechanism to achieve local and global attention while maintaining efficient computation in figure 5. The block starts with the input of the Residual Block followed by the Window Partition operation which partitions the input data into non-overlapping windows. The MHSA is applied in each segment to capture complex relationships, after which the outputs of nearby windows are merged by Window Merging to enrich the context. A Dropout Layer is used for regularization of the merged output and a skip connection (“+”) keeps the original input for residual learning. There is also another Layer Normalization and another Multi-Layer Perceptron (MLP) to further refine the features, and an extra Dropout Layer to provide additional regularization. The final skip connection combines the output from the MLP with the normalized input after the attention layer to form a complete output, which will contain information from various scales, to be passed to the rest of the network.

**Figure 5:**
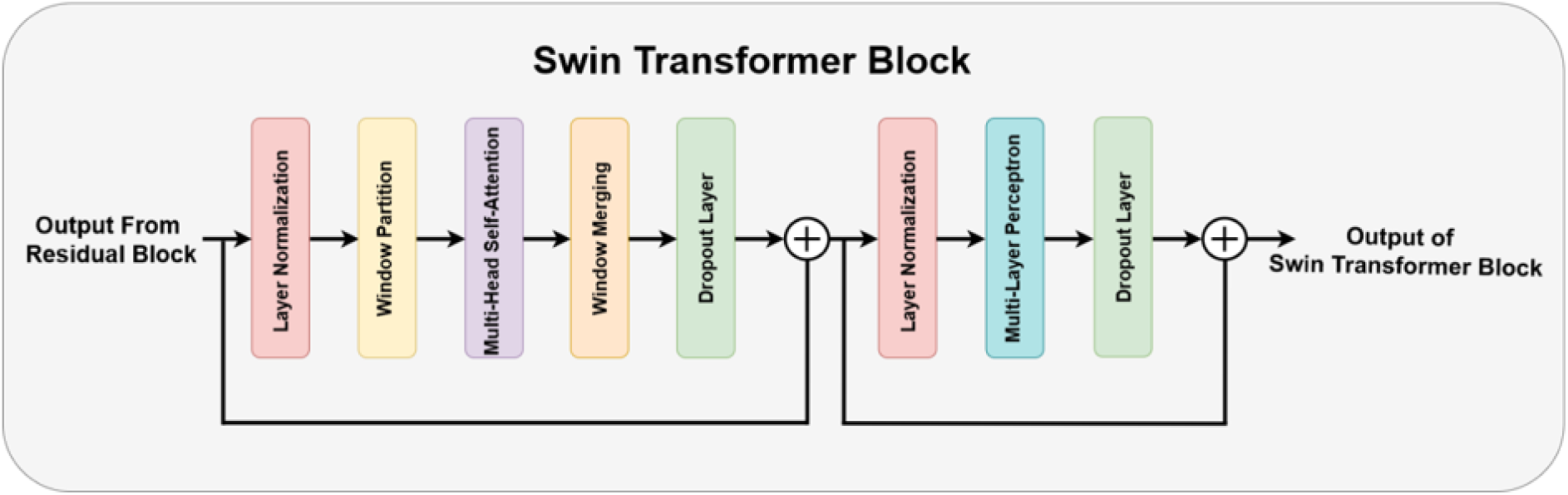
Swin Transformer Block.

### Equations and Operations

#### Layer Normalization

The input from the residual block is normalized before proceeding to the next step. Mathematically, this operation is represented in equation 3, Where *X* is the input, and *X_norm_* is the normalized output.

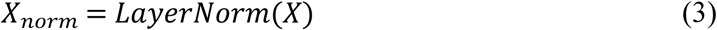

#### Window Partitioning

The input feature map is partitioned into smaller windows of size window_size X window_size, making the process computationally efficient; it is given in equation 4, Where *X_ω_* represents the windows formed from the input.

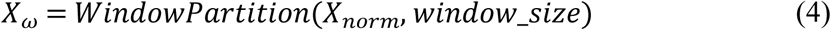

#### MHSA

Attention is applied within each window to capture relationships between different tokens inside that window. This is represented by equation 5, Where *A_ω_* is the attention output within each window, and MHSA computes the self-attention.

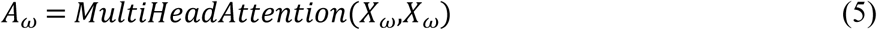

#### Window Merging

After applying self-attention, the windows are merged back into the original spatial dimensions, as shown in equation 6, Where *X_merge_* is the merged output from the attention blocks.

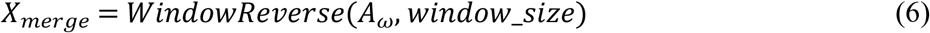

#### Residual Connection after Attention

is added after merging, incorporating the original input to the block, ensuring the model can learn identity mappings more efficiently, as shown in equation 7, Where *X* is the original input from the previous block and *X*′ is the output after adding the residual connection.

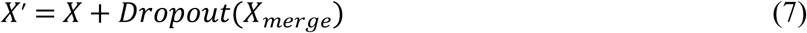

#### Layer Normalization Before MLP

Another layer normalization is applied before passing the result into the Multi-Layer Perceptron (MLP) block, as shown in equation 8:

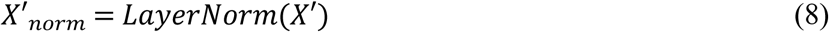

#### Feed-Forward Network (MLP)

*The* normalized output is passed through a MLP containing of two fully connected layers. This helps the network capture more complex patterns. Mathematically, this is represented in equation 9:

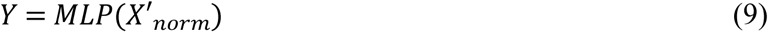

#### Residual Connection after MLP

Finally, another residual connection is added after the MLP block, again ensuring the gradients can flow efficiently through the model, as shown in equation 10, Where *Z* is the final output of the Swin Transformer Block, which will be passed on to the next stage in the network.

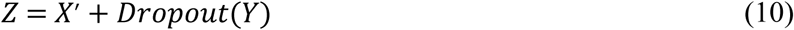

### Attention Mechanisms

It’s improves the model’s ability to focus on important parts of the input, essential for the correct segmentation of images. The block receives inputs from the Decoder and the Swin Transformer Block as shown in figure 6. Input is first passed through individual 2D Convolutional layers to extract different features of the inputs. The processed outputs are then added together to allow important features to be added from both pathways. This combined feature map then goes through a ReLU Activation Block and another 2D Convolutional block, to refine the features further. The next Sigmoid Activation Block produces attention weights which guide the model to focus more on features of the image that are more pertinent. An up-sampling operation is then applied to rescale the feature map to the desired spatial dimensions and a final 2D Convolutional layer is used to refine the depth of the output for the final segmentation task. This kind of operations enables the model to dynamically focus on the most relevant features and improve the accuracy of the segmentation output.

**Figure 6:**
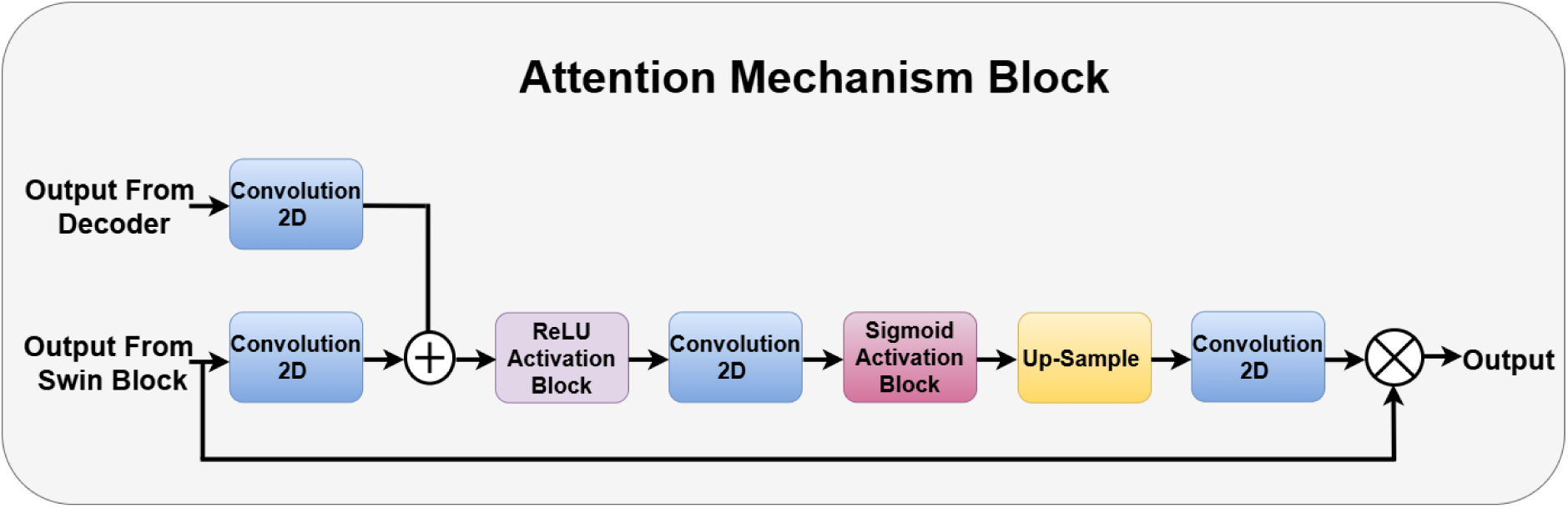
Attention Mechanism Block.

### Equations and Operations

Convolution Operation from Decoder and Swin Block are shown in equations 11 and 12, Where: *D_conv_* and *S_conv_* are the outputs of 2D convolution applied to the decoder output *D* and the Swin block output *S*, respectively. *W_d_* and *W_s_* represent the convolution weights. *b_d_* and *b_s_* are the biases and ∗ denotes the convolution operation.

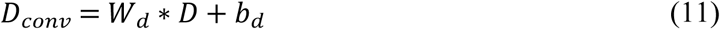

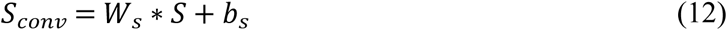

Element-wise Addition (Residual Connection) is given in equation 13, Where *F_add_* is the sum of the convolved outputs.

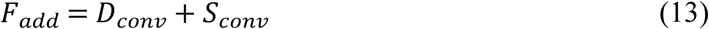

The ReLU is presented in equation 14, Where *F_ReLU_* is the result after applying the ReLU activation function.

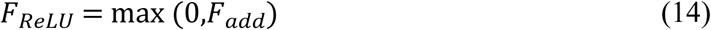

The second Convolution equation is presented in equation 15, Where *F_Conv_*_2_ is the output of the second 2D convolution layer applied to the ReLU output, and *W*_2_ and *b*_2_ are the weights and biases for the second convolution.

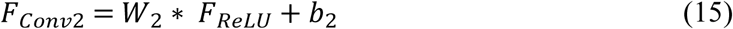

Sigmoid Activation (Attention Generation) is given in equation 16, Where *F_sigmoid_* is the attention map generated by the sigmoid activation function.

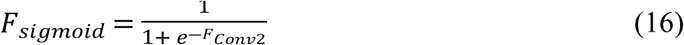

Up-sampling Operation is presented in equation 17, Where *F_upsample_* is the up-sampled result to match the desired resolution.

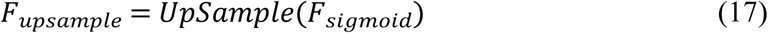

The final Convolution is given in equation 18, Where *Output* is the final output of the attention mechanism block, and *W*_3_ and *b*_3_ are the weights and biases for the final convolution layer.

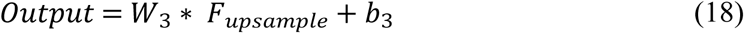

### Proposed RSAUNet Model

The code is based on a U-Net model with Swin Transformer blocks, residual convolutional blocks, and attention mechanisms to achieve improved image segmentation. The “window_partition” and “window_reverse” functions segment the input images into smaller patches and reassemble them, which is crucial for the window-based self-attention mechanism in the Swin Transformer. The “SwinTransformerBlock” class represents a single Swin Transformer block, combining MHSA for local window attention with a feed-forward neural network (FFN) to effectively capture spatial hierarchies. Residual connections preserve input features through several layers to enhance the stability and convergence in model training. The “residual_conv_block” function defines a convolutional block with residual connections, allowing the model to learn more complex patterns and helping to prevent the vanishing gradient problem. The “attention_block” function introduces an attention mechanism into the model, enabling it to focus on the most relevant features of the input image, which improves the accuracy of segmentation.

The create_unet_model function is used to set up the architecture of the U-Net with Swin Transformer blocks added at every level of the encoder and decoder to capture the local and global context of the image. In the encoder section, the convolution layers are combined with the Swin Transformer to down-sample and extract features. Meanwhile, the decoder conducts upsampling with the aid of attention mechanisms to accurately reconstruct the segmented output, as illustrated in figure 7. The hybrid model works effectively by using the convolutional and transformer-based approaches in dealing with complex segmentation problems, like medical images, where boundary detection is crucial. Model compilation and summary the model is now ready to train on image data. It is particularly suited to high-resolution applications like outlining regions of interest, such as delineating lesions or anatomical structures within MRI scans.

**Figure 7:**
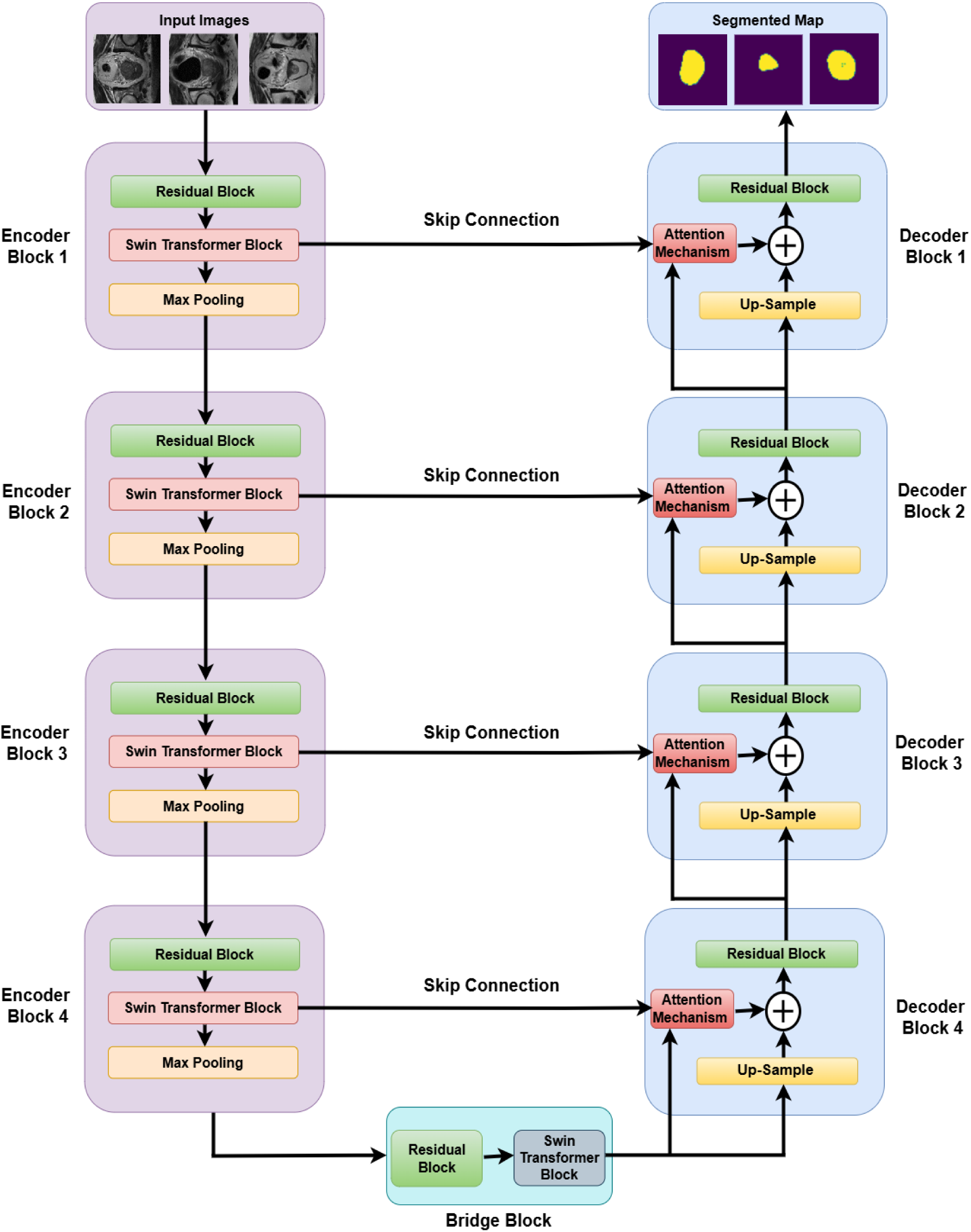
Proposed RSAUNet Model.

**Algorithm for Residual Attention Swin U-Net (RSAUNet) Model**

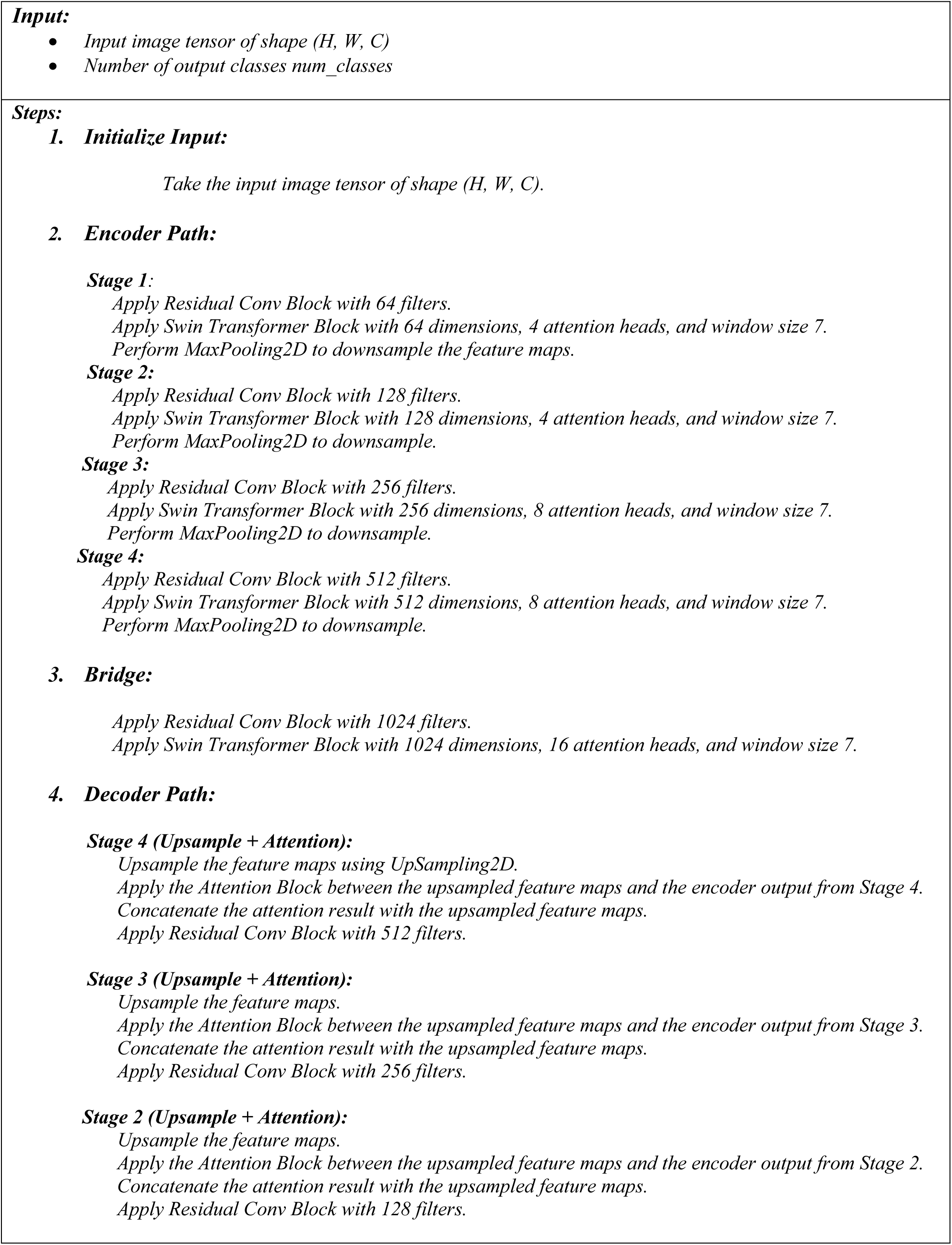
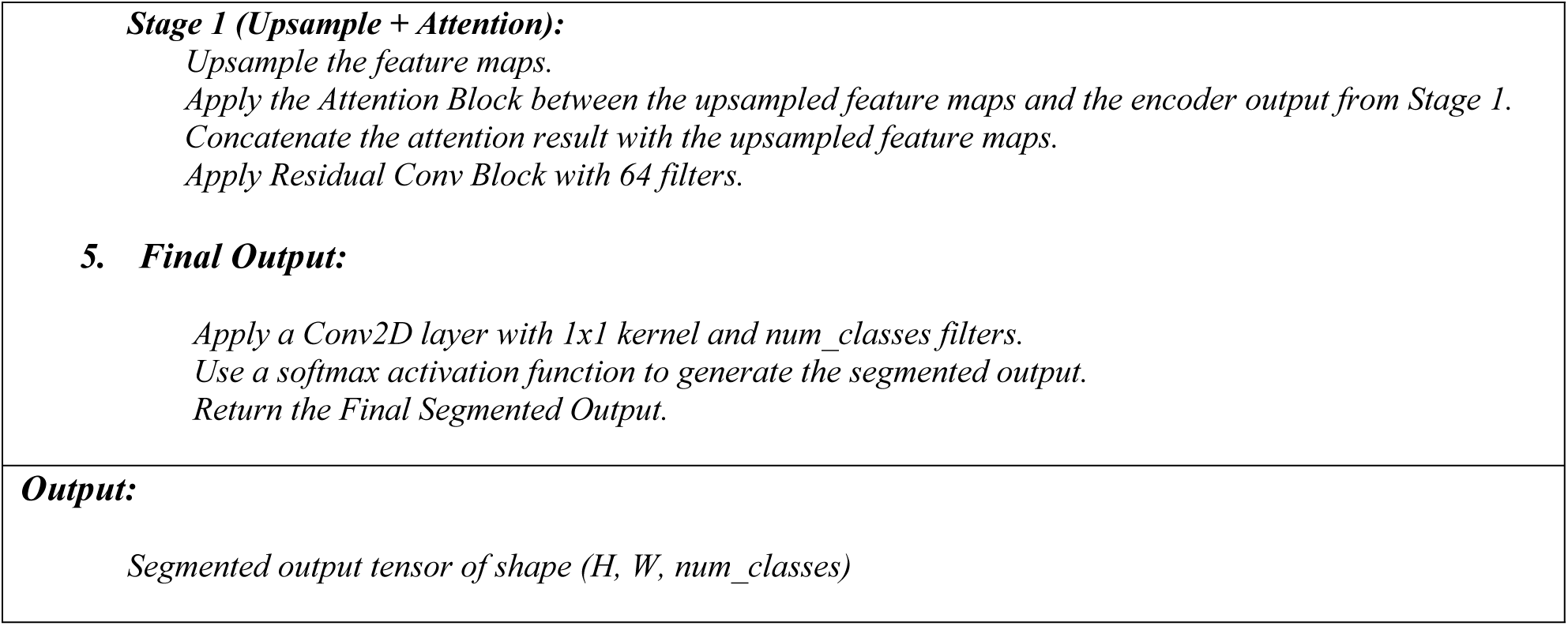

## Results and Discussion

The Results section is organized to evaluate the model’s performance for different configurations and improvements. The analysis begins by analyzing the baseline U-Net model, then analyzes the performance improvement of adding Residual Blocks, Swin Transformer Blocks and Attention Mechanisms. The assessment for each configuration is based on training and validation loss, accuracy, DC and Mean Intersection over Union (M IoU). By combining all these improvements in the proposed RSAUNet model, it has demonstrated outstanding results in all the metrics. The findings then explore cutting-edge segmentation strategies and provide a detailed comparison to highlight improvements in training stability, segmentation performance and adaptability to unseen data.

### Results on UNet

The learning progress of the U-Net model is shown visually in 4 subplots (a-d) in figure 8, over a total of 300 epochs. The Loss curves (Figure 8(a)) reveal that losses for training and validation sets both converge quickly to low values of 0.02 and 0.03, respectively, confirming good learning and lack of overfitting. The Accuracy is shown in figure 8(b) and both curves converge to a near perfect accuracy at the end (0.98 and 0.971 respectively). The Training and Validation DC are two important metrics to assess the quality of the segmentation and they are both 0.98 for training and validation respectively (see Figure 8(c)). Finally, in Figure 8(d), the increasing trend in both the Training and Validation Mean IoU is observed, with the Training IoU at 0.928 and the Validation IoU stabilizing around 0.917.

**Figure 8:**
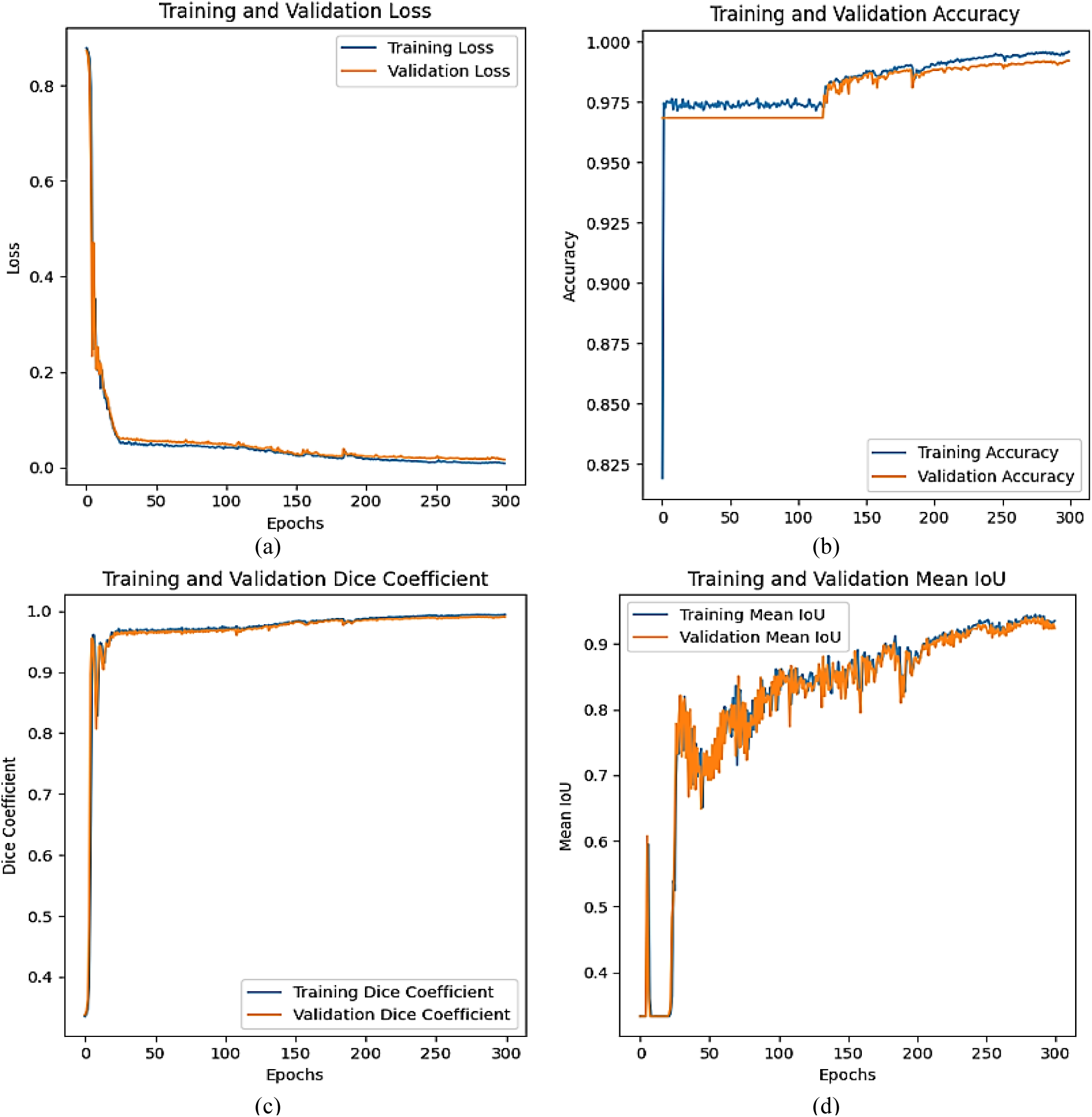
Graphical Analysis of UNet Model Training and Validation (a) Loss, (b) Accuracy, (c) Dice Coefficient & (d) Mean IoU.

The final learning outcomes of the U-Net model after being trained on a segmentation task are presented in Table 2 and Figure 9. The model had negligible overfitting with Loss values of 0.02 and 0.03. The Training Accuracy achieved was 0.98 while the Validation Accuracy was slightly lower at 0.971, indicating that the model was successfully trained and it also performed well on the validation set. The Dice Coefficients were both 0.98 which shows a high accuracy rate for the segmentation tasks. Moreover, the Training MIoU was 0.928, and the Validation Mean IoU was 0.917.

**Figure 9:**
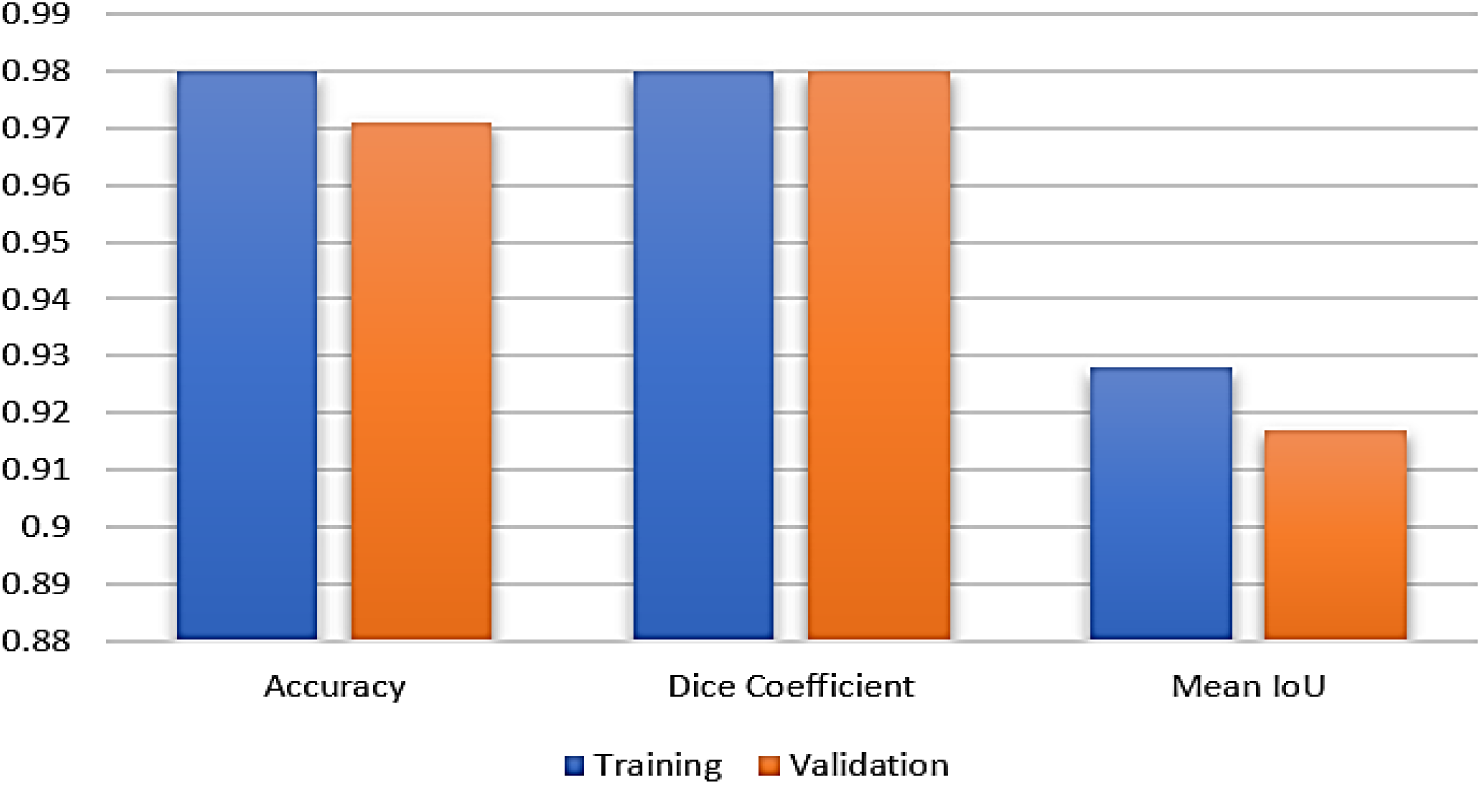
Training and Validation Analysis of UNet.

**Table 2:** Final Learning Results of UNet.

| Parameters | Training | Validation |
| --- | --- | --- |
| Loss | 0.02 | 0.03 |
| Accuracy | 0.98 | 0.971 |
| Dice Coefficient | 0.98 | 0.98 |
| Mean IoU | 0.928 | 0.917 |

### Results on UNet + Residual Block

The learning dynamics for the U-Net model (with Residual Block) are presented in Figure 10 in four subplots, (a) through (d). The Loss are shown in Figure 10(a), with a significant decrease in loss values to 0.01, and to 0.04. Figure 10(b) shows the Training and Validation Accuracy, which quickly reaches 0.981 and validation accuracy is around 0.93 with good generalization. Figure 10(c) shows that the Training and Validation Dice Coefficient are both high, reaching a peak of close to 0.99. Lastly, Figure 10(d) shows the Mean IoU, with both metrics showing steady improvement; the training and the validation IoU settles at around 0.916 and 0.911.

**Figure 10:**
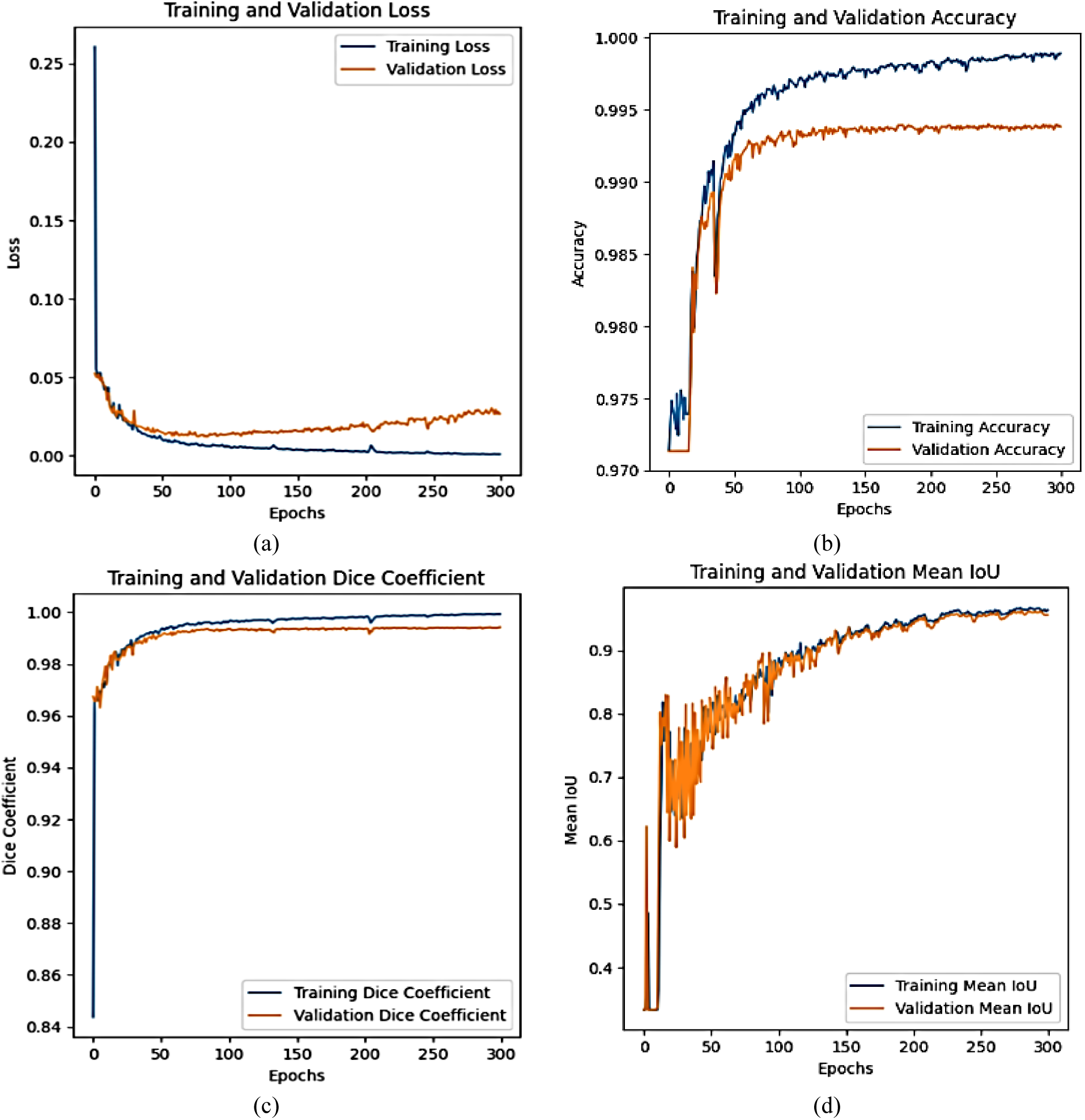
Graphical Analysis of UNet + Residual Training and Validation (a) Loss, (b) Accuracy, (c) Dice Coefficient & (d) Mean IoU.

Table 3 and Figure 11 display the learning outcome of the U-Net model with the Residual Block and the effect it induces on the model performance metrics. There was no overfitting as the model performed with Loss of 0.01 and 0.04, suggesting that the model learned well, with slight complexity in the validation data. The accuracy of training is 0.981 and that of validation is 0.93 which shows good accuracy with a slight drop during validation. The Training DC was 0.994 and the Validation DC was 0.991, indicating the model’s ability to achieve high precision even when tested on new data. Moreover, the mean IoU of the training and validation sets are 0.916 and 0.911.

**Figure 11:**
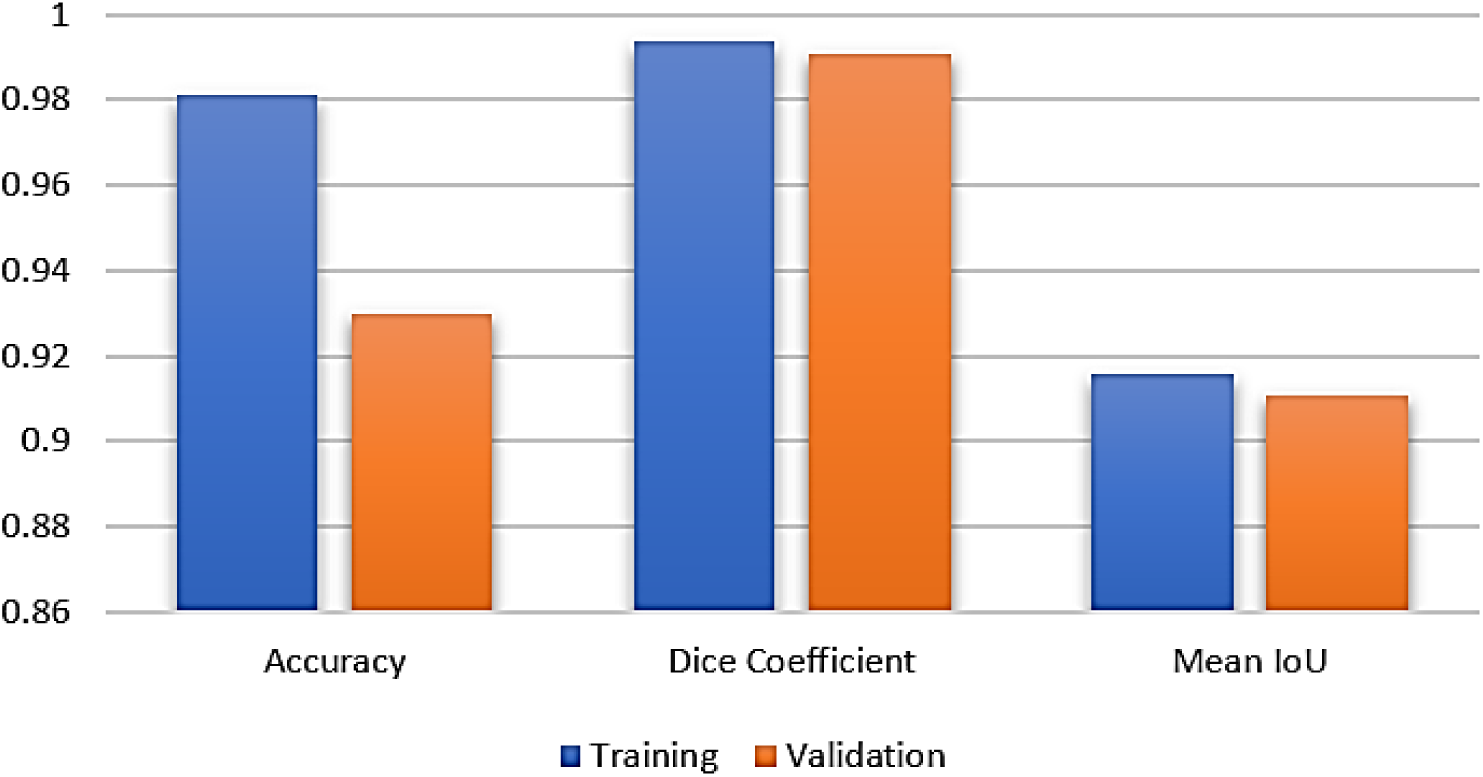
Training and Validation Analysis of UNet + Residual Block

**Table 3:** Final Learning Results of UNet + Residual Block.

| Parameters | Training | Validation |
| --- | --- | --- |
| Loss | 0.01 | 0.04 |
| Accuracy | 0.981 | 0.93 |
| Dice Coefficient | 0.994 | 0.991 |
| Mean IoU | 0.916 | 0.911 |

### Results on UNet + Residual + Swin Transformer Block

The learning dynamics of the U-Net model featuring the combined Residual and Swin Transformer Blocks are shown in four subplots (a) to (d) in Figure 12. The Loss curves are shown in figure 12(a) and both curves show meaningful convergence around 0.02 and 0.03, respectively, implying that the model has converged well and there is not much overfitting. Figure 12(b) depicts the Training and Validation Accuracy, with good accuracies of 0.982 and 0.973. The Training and Validation Dice Coefficient are shown in figure 12(c), both curves attained 0.981. Finally, Figure 12(d) shows the Mean IoU, both of which show an increasing trend; the Training IoU are 0.919 and the Validation IoU are 0.912.

**Figure 12:**
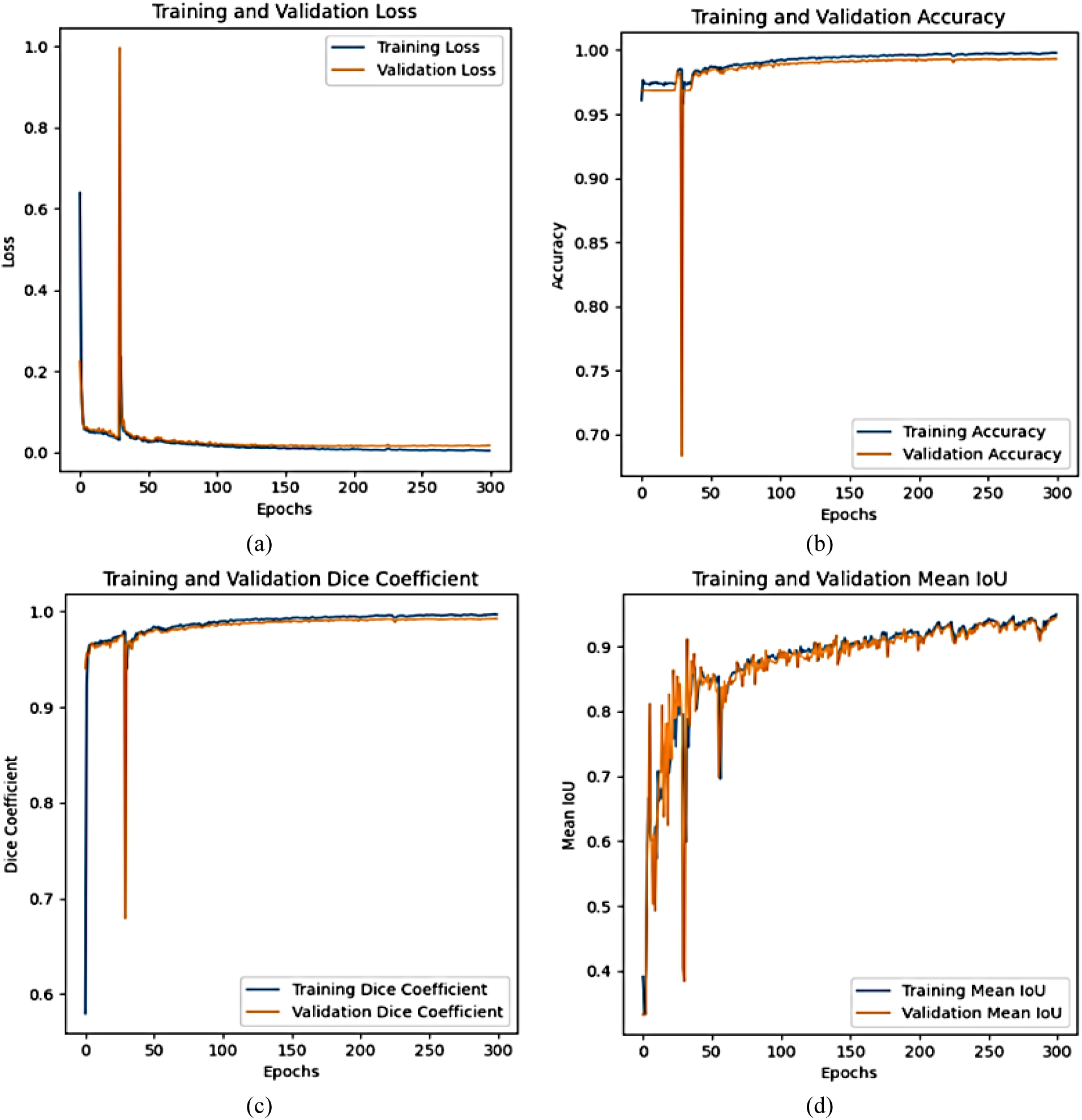
Graphical Analysis of UNet + Residual + Swin Transformer Block Training and Validation (a) Loss, (b) Accuracy, (c) Dice Coefficient & (d) Mean IoU.

Table 4 and Figure 13 show the final learning results of U-Net model that utilizes both Residual and Swin Transformer Blocks, which is an advanced architectural configuration for improving the effectiveness of the segmentation. The Losses are 0.02 and 0.03 indicate that the model’s training was stable and efficient, with little evidence of overfitting. The accuracy was 0.982 and 0.973. The DC was 0.981 and 0.972. The Training and Validation Mean IoU were also 0.919 and 0.912.

**Figure 13:**
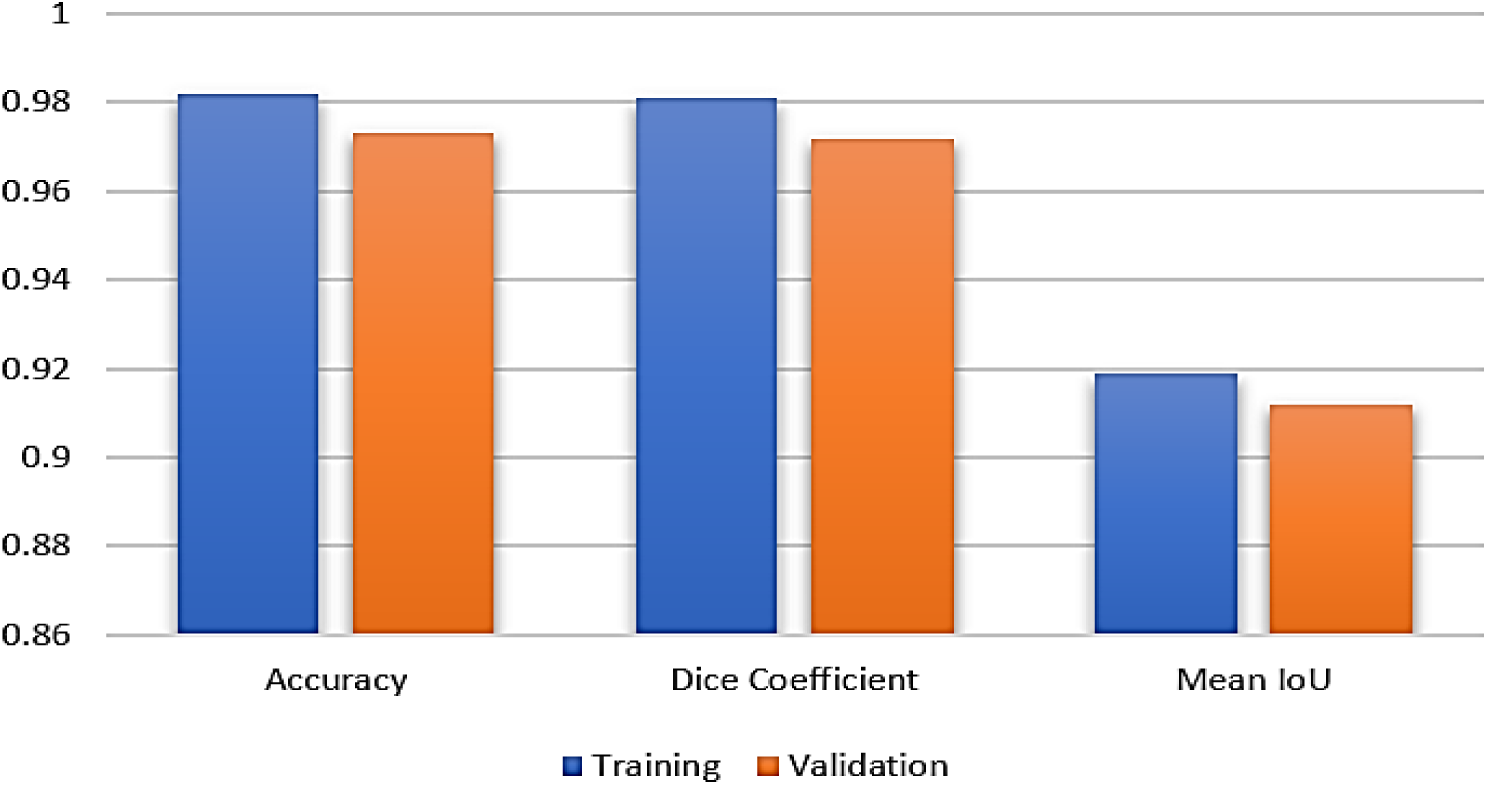
Training and Validation Analysis of UNet + Residual + Swin Transformer Blocks.

**Table 4:** Final Learning Results of UNet + Residual + Swin Transformer Block.

| Parameters | Training | Validation |
| --- | --- | --- |
| Loss | 0.02 | 0.03 |
| Accuracy | 0.982 | 0.973 |
| Dice Coefficient | 0.981 | 0.972 |
| Mean IoU | 0.919 | 0.912 |

### Results on UNet + Residual + Swin Transformer Block + Attention Mechanisms Block

The performance of the model is shown in Figure 14, which is subdivided into four subplots, (a), (b), (c), and (d). The Loss curves shown in Figure 14(a) both declines rapidly and converge to low values (0.01 for training loss and 0.03 for validation loss), indicating good learning and low chances of overfitting. It is observed from Figure 14(b) that Training and Validation Accuracy are high, with the highest accuracy of 0.995 on the Training set and 0.993 on the Validation set, reflecting the model’s strong generalisation capability. The DC is shown in figure 14(c), and is similarly very high, with a training DC of 0.983 and a validation DC of 0.971. Lastly, Figure 14(d) depicts the Training and Validation Mean IoU, which both increase steadily to 0.939 and 0.934, respectively.

**Figure 14:**
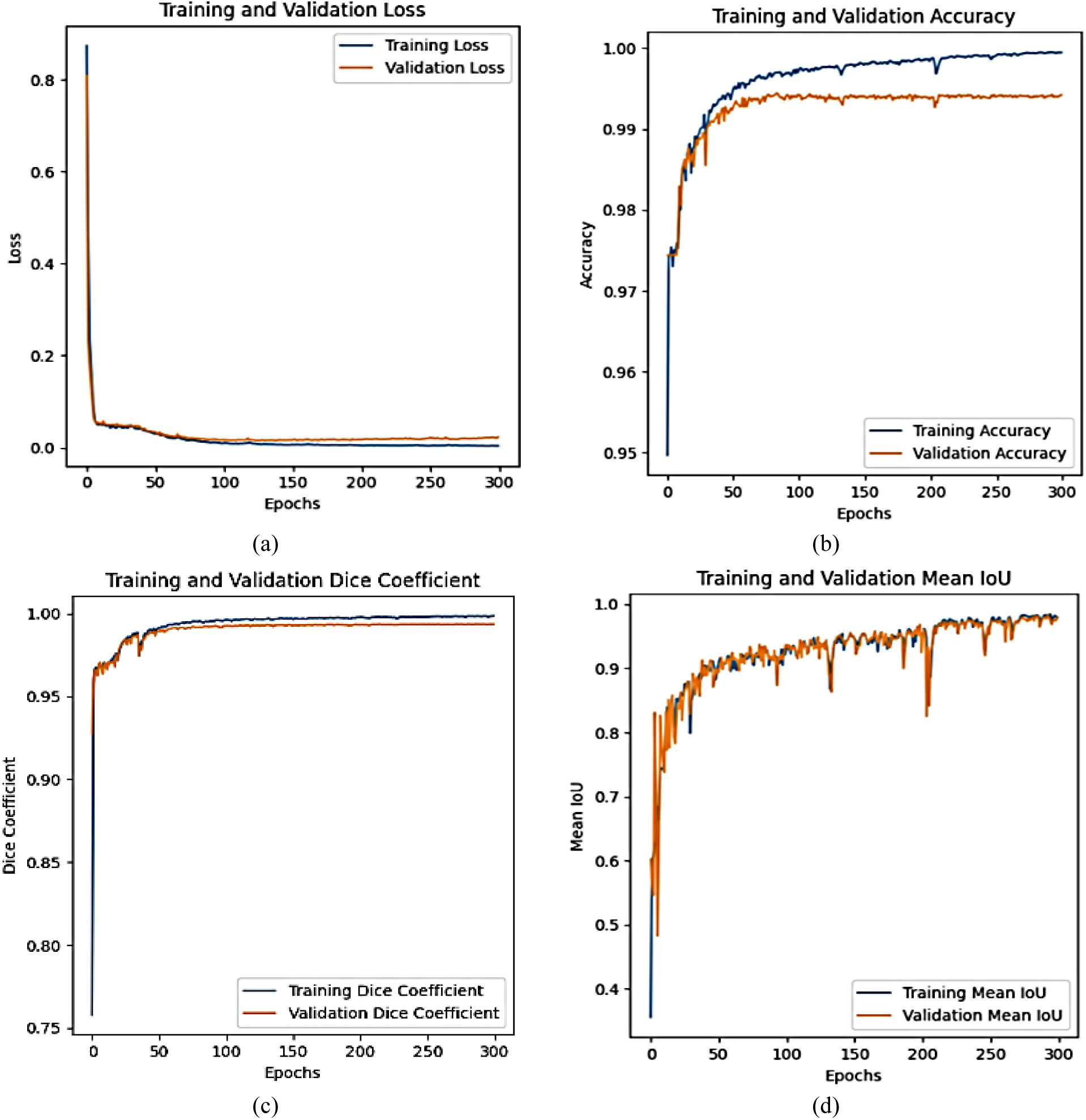
Graphical Analysis of UNet + Residual + Swin Transformer Block + Attention Mechanisms Block Training and Validation (a) Loss, (b) Accuracy, (c) Dice Coefficient & (d) Mean IoU.

Table 5 and Figure 15 show the final learning outcomes for the U-Net model with Residual, Swin Transformer and Attention Mechanisms Blocks. The aim of this model configuration is to take the best of each of the three components to achieve better performance in segmentation tasks. It has a Training and validation loss of 0.01 and 0.03, which are extremely low, suggesting very good convergence and good generalization capabilities. On unseen data, Accuracy was 0.995 and 0.993, signifying excellent performance on previously unseen data. Training DC was 0.983, and the Validation DC was nearly 0.971. The Training Mean IoU and the Validation Mean IoU were large, 0.939 and 0.934, respectively.

**Figure 15:**
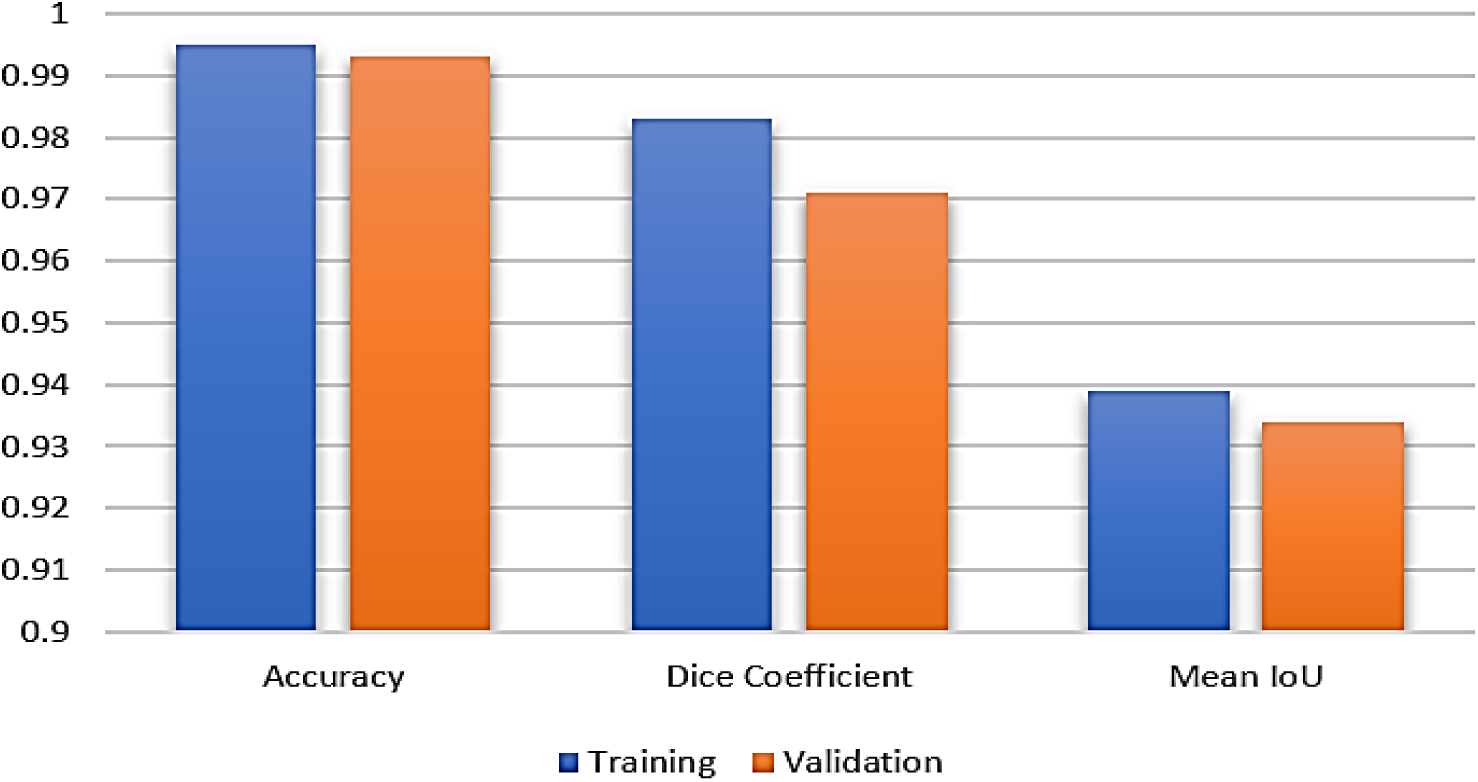
Training and Validation Analysis of UNet + Residual + Swin Transformer + Attention Mechanisms Block.

**Table 5:** Final Learning Results of UNet + Residual + Swin Transformer Block + Attention Mechanisms Block.

| Parameters | Training | Validation |
| --- | --- | --- |
| Loss | 0.01 | 0.03 |
| Accuracy | 0.995 | 0.993 |
| Dice Coefficient | 0.983 | 0.971 |
| Mean IoU | 0.939 | 0.934 |

### Results on Proposed RSAUNet Model

Figure 16 presents a visual analysis of the training and test of the model of RSAUNet in the four graphs marked (a), (b), (c) and (d). Efficient learning is illustrated by the Loss curves in Figure 16(a), which rapidly decrease to very low levels (0.01 for training, 0.02 for validation). The model’s performance in segmentation tasks is reliable as confirmed by the Training and Validation Accuracy from Figure 16(b) both curves are high with training accuracy of 0.999 and validation accuracy of 0.998. Both the Training and Validation Dice Coefficient values are low in all cases shown in figure 16(c) signifying that the model has a good performance in accurately determining different regions during segmentation process. Lastly, Figure 16(d) presents the Training and Validation Mean IoU; the metrics demonstrate a gradual rise, reaching 0.979 for Training IoU and 0.965 for Validation IoU, which further highlights the model’s competence in producing accurate segmentations even with varied data inputs.

**Figure 16:**
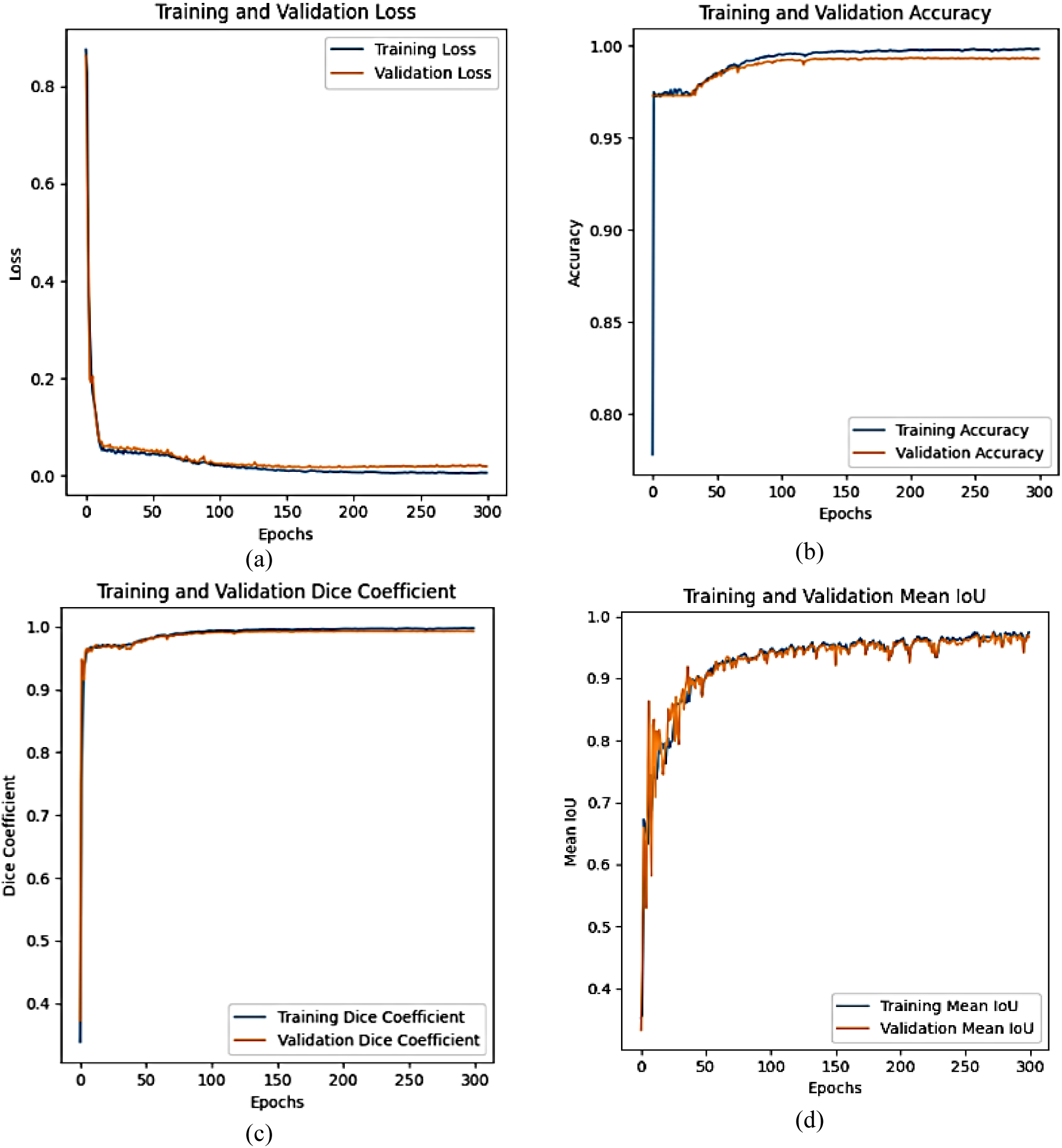
Graphical Analysis of Proposed RSAUNet Model Training and Validation (a) Loss, (b) Accuracy, (c) Dice Coefficient & (d) Mean IoU.

The proposed model, RSAUNet, has demonstrated its ability for segmentation tasks in the final learning outcome as shown in Table 6 and Figure 17. The loss of the proposed RSAUNet model resulted in a very low value (0.01 and 0.02), which shows that the model was trained very well. The Accuracy is an impressive 0.999 and 0.998 indicating good generalization to new unseen data. The DC are 0.999 and 0.998. The Mean IoU are 0.979 and 0.965, which indicates good intersection among the segmentation masks predicted and actual segmentation masks.

**Figure 17:**
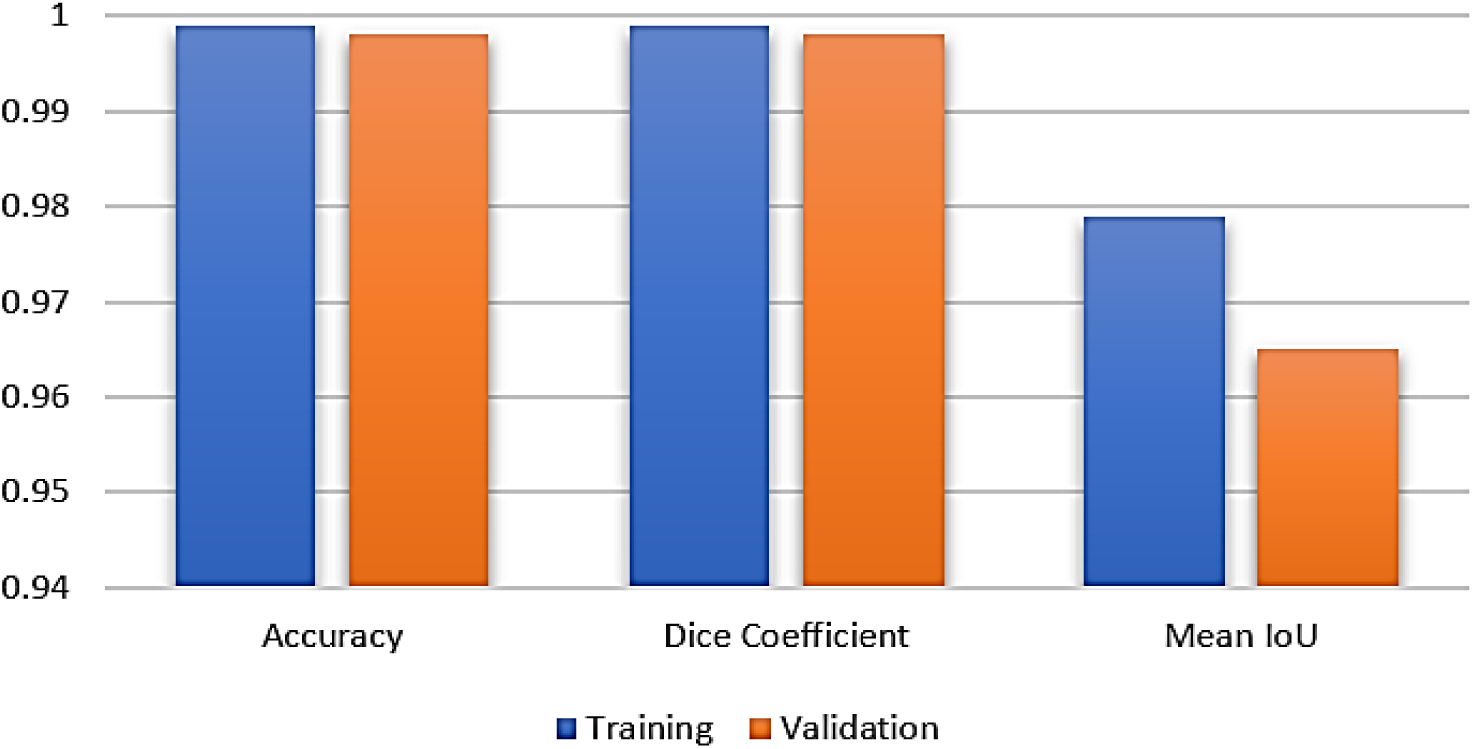
Training and Validation Analysis of Proposed RSAUNet.

**Table 6:** Final Learning Results of Proposed RSAUNet.

| Parameters | Training | Validation |
| --- | --- | --- |
| Loss | 0.01 | 0.02 |
| Accuracy | 0.999 | 0.998 |
| Dice Coefficient | 0.999 | 0.998 |
| Mean IoU | 0.979 | 0.965 |

### Ablation Study

The improvements of the model with different architectural changes is presented in table 7. When adding the Residual Blocks to the baseline U-Net with accuracy of 0.97 and Dice coefficient of 0.98, the validation accuracy decreases slightly to 0.93, while the Dice coefficient increases to 0.99, which means that the Residual Blocks can capture more complex features. The residual-enhanced U-Net with Swin Transformer blocks has comparable training and validation metrics with a marginal decrease in the Dice coefficient and IoU. The addition of attention mechanisms and Swin Transformer blocks is however enough to yield dramatic performance gains, achieving 1.00 training accuracy, 0.99 validation accuracy, and also a rise in the Dice coefficient (0.97) and IoU (0.93). Lastly, the proposed model, called RSAUNet, that combines all these components achieves the best overall balance of metrics, with a validation accuracy of 0.98, DC of 0.99, and a mean IoU of 0.96, demonstrating the best generalisation capability and ability to segment prostate tissues accurately.

**Table 7:**
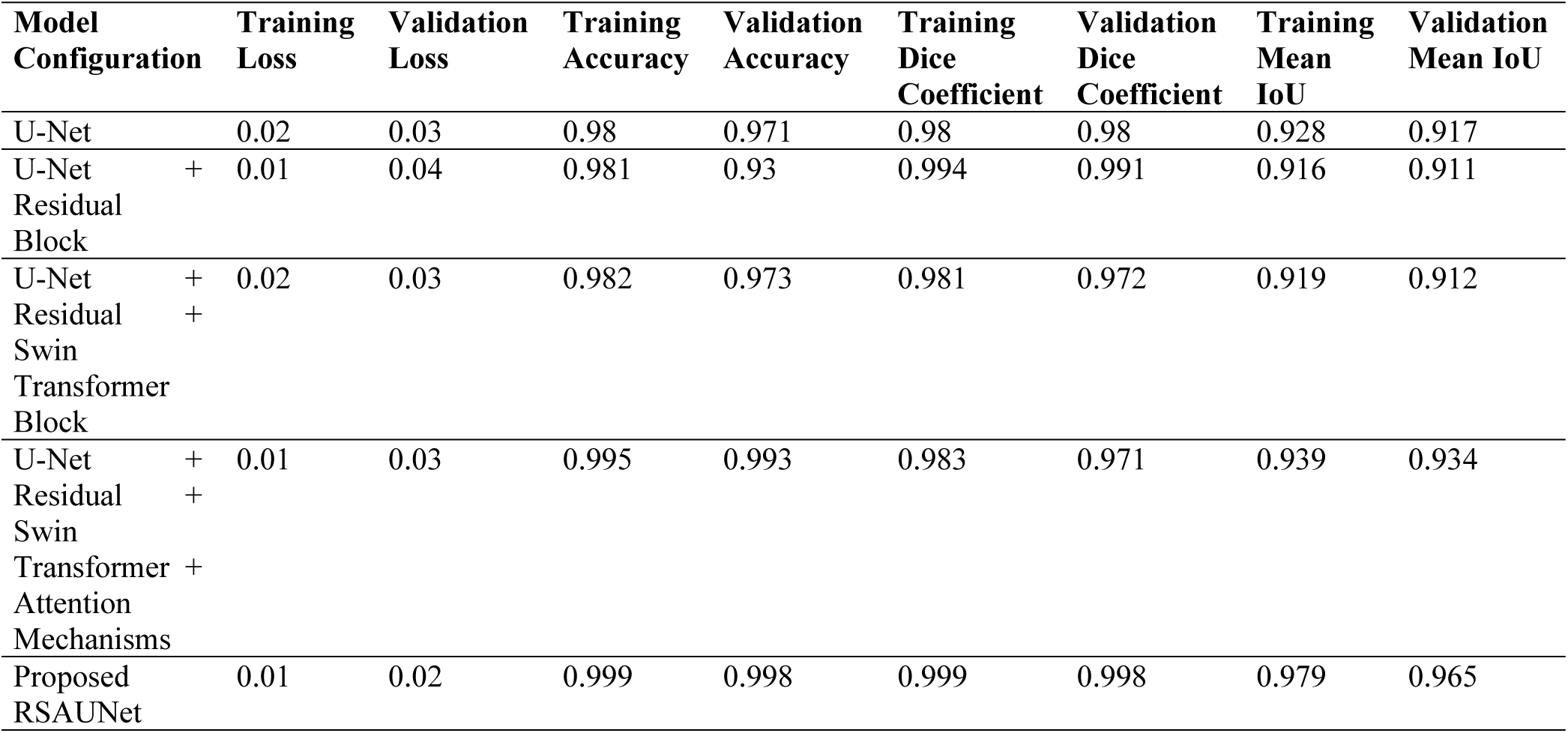
Ablation Study.

The loss analysis for various model configurations is presented in Figure 18, highlighting the impact of each architectural improvement on the model’s performance. The bar graph compares their training loss (blue bars) and validation loss (orange bars). By adopting residual blocks, Swin Transformer blocks, and attention mechanisms, the overall performance is enhanced, resulting in a drop in training and validation losses. The U-Net with residual blocks has the highest validation loss, and the Proposed RSAUNet has the lowest training and validation losses, suggesting that it works well in learning and generalizing to the validation set. The visual comparison shows that each of the architectural additions leads to better performance of the model, with the best one finding the minimum difference between the loss on the training and validation set, which is the RSAUNet model.

**Figure 18:**
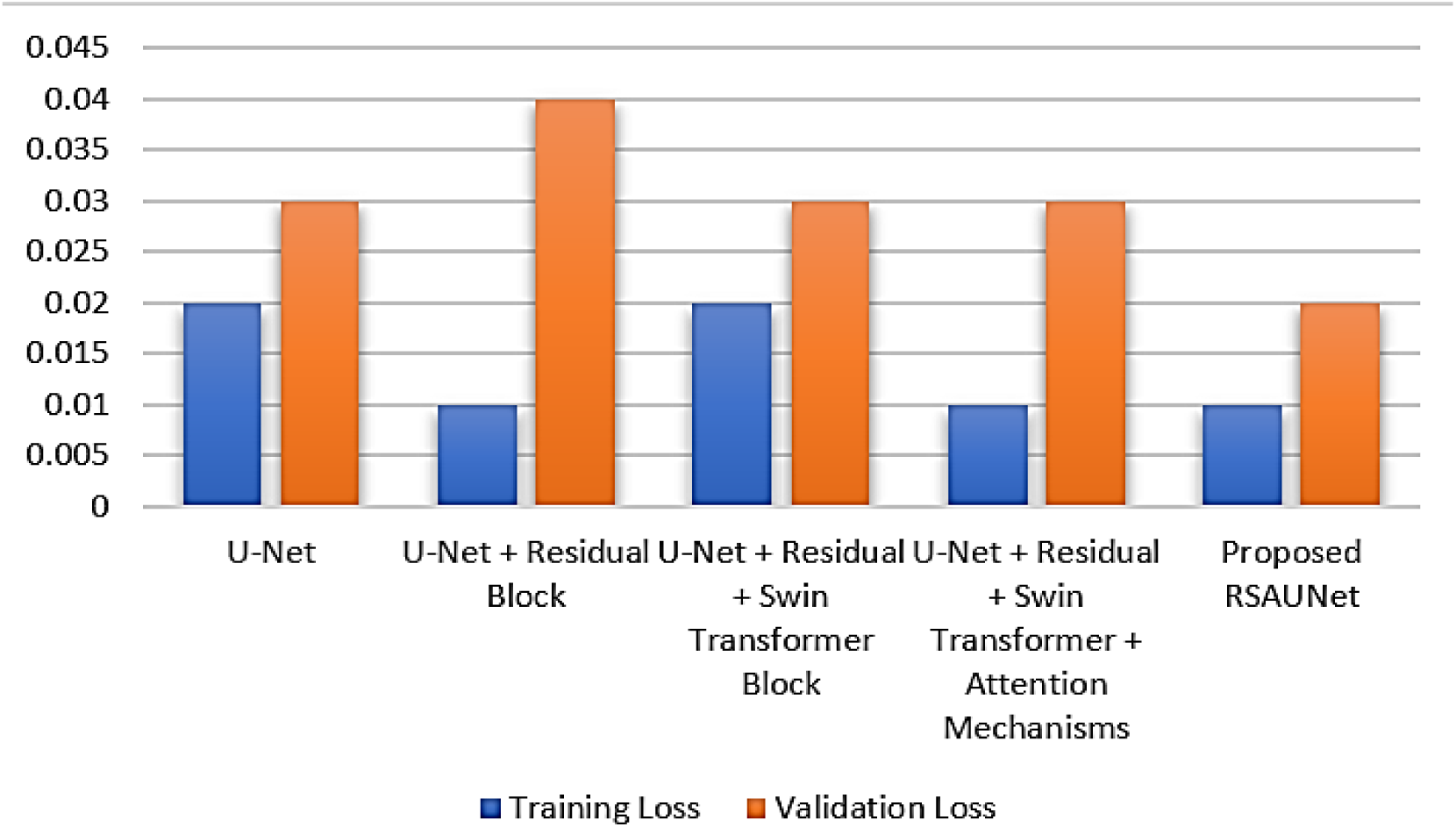
Loss Analysis.

The accuracy of various model configurations is compared in figure 19. The U-Net and U-Net + Residual models are performing well in terms of training accuracy, but have a slight loss in validation accuracy, signifying overfitting. The Swin Transformer and attention mechanisms are added and the validation accuracy increases considerably, with the Swin Transformer + Residual + U-Net + Attention Mechanisms model achieving almost training and validation accuracy of 0.995 and 0.993. The Proposed RSAUNet shows a good training and validation accuracy of 0.999 and 0.988, which highlights a good generalisation to unseen data.

**Figure 19:**
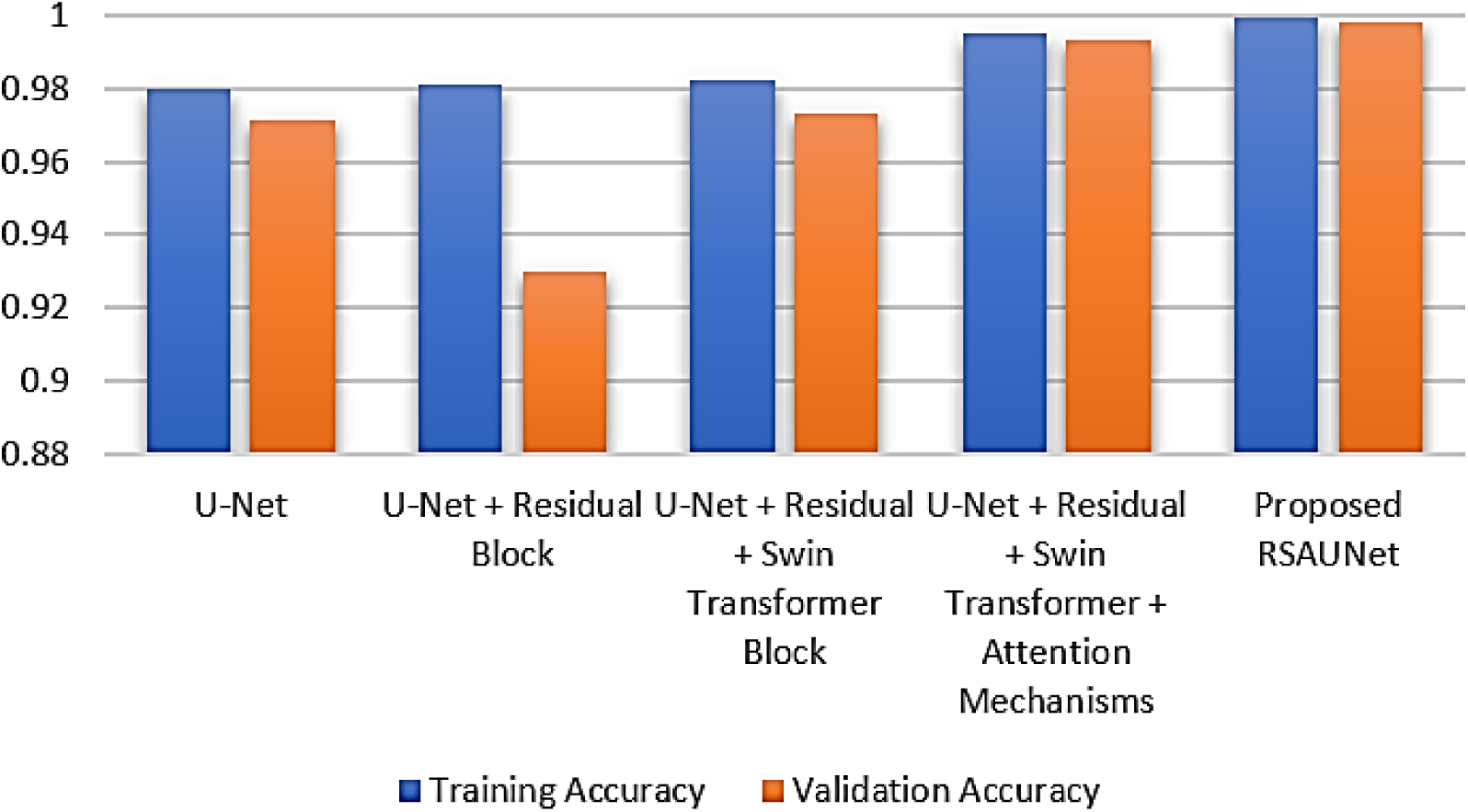
Accuracy Analysis.

The analysis of DC is represented by figure 20 in which different configurations of the model are represented. The models U-Net and U-Net + Residual models achieve high Dice Coefficient with U-Net + Residual Block achieving near perfect training and validation results. Validation Dice Coefficient degrade slightly as Swin Transformer and attention mechanisms are added on while the value of training is high. For the Proposed RSAUNet, the highest DC is obtained in the training (0.999) and validation (0.998) phase, indicating an excellent balance.

**Figure 20:**
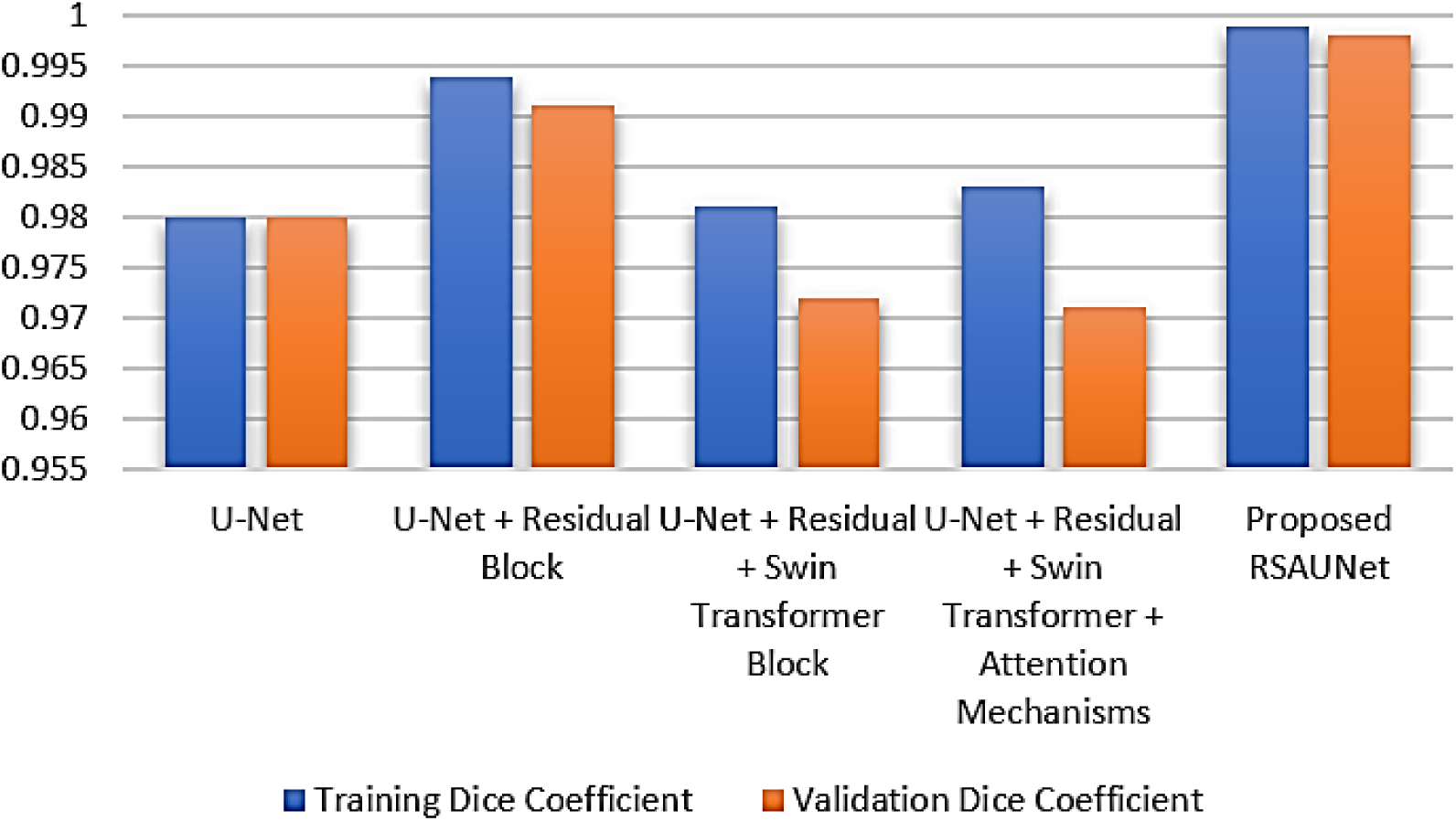
Dice Coefficient Analysis.

The results of Mean IoU analysis of the models are presented in figure 21. The blue bars are the Training Mean IoU and orange bars are the Validation Mean IoU. The U-Net model has a moderate performance as it has a slightly lower IoU when evaluated on the Validation Data than when assessed on the Training Data. The IoU values change slightly as the Residual Blocks, Swin Transformer Blocks and Attention Mechanisms are introduced. Incorporating the Swin Transformer Block does not result in significant gains over the previous models in terms of IoU, but the Proposed RSAUNet achieves a remarkable gain in both Training and Validation Mean IoUs, and the best performance across all the configurations.

**Figure 21:**
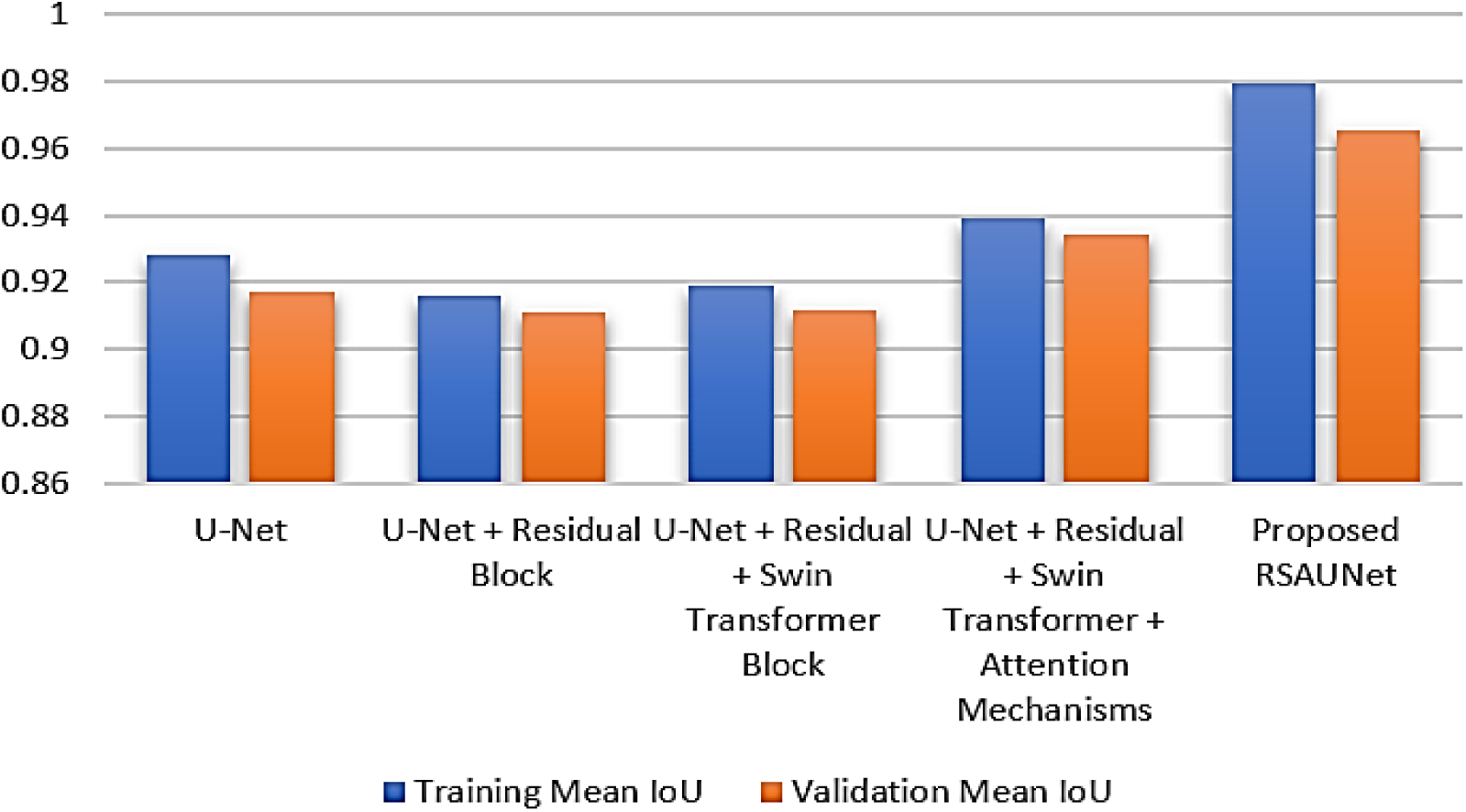
Mean IoU Analysis.

### Visual Analysis

To see the comprehensive investigation of the segmentation performance of the prostate across various model configurations, refer to Figure 22, which shows the U-Net to Proposed RSAUNet model. Results are shown in the second column (the true MRI masks manually annotated by the experts), and in the other columns (the predicted masks produced by each model configuration). Yellow regions in both masks (true and predicted) represent segmented prostate regions. The predicted masks are in good agreement with the actual masks, and as observed. The comparison reveals the benefits of the modifications realized with the various versions, and the results demonstrate the gains in its segmentation accuracy and consistency in moving from U-Net to the proposed RSAUNet. Residual blocks, Swin Transformer blocks, and attention mechanisms are used to further improve the segmentation, and the RSAUNet model obtains the most accurate and reliable segmentation results for the prostate.

**Figure 22:**
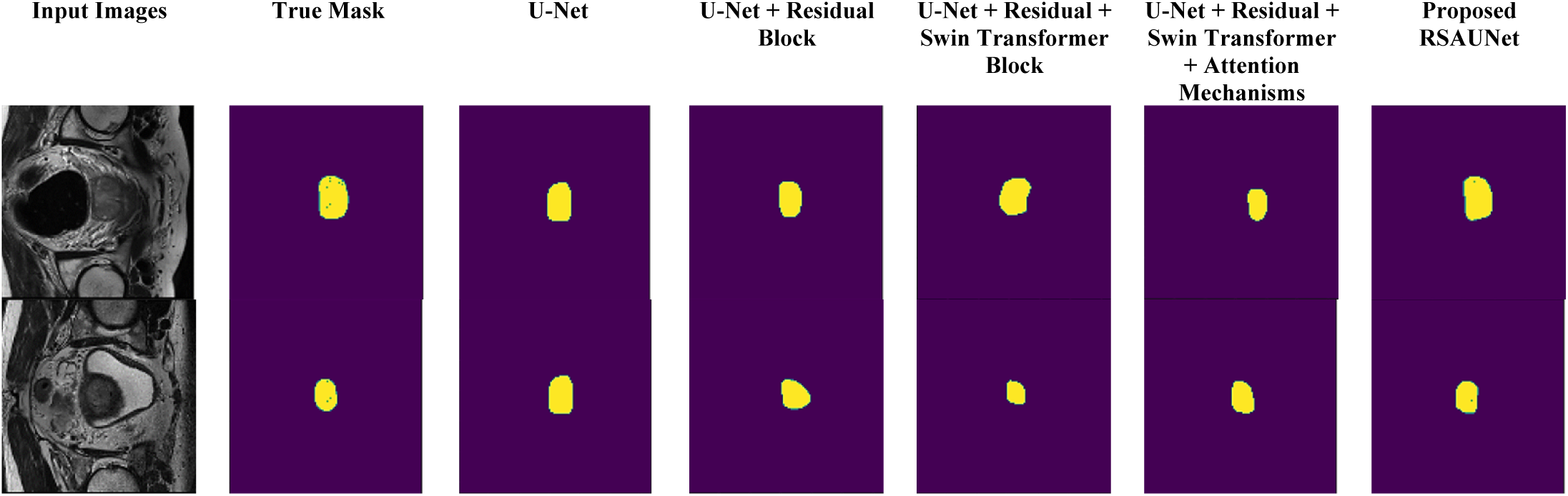
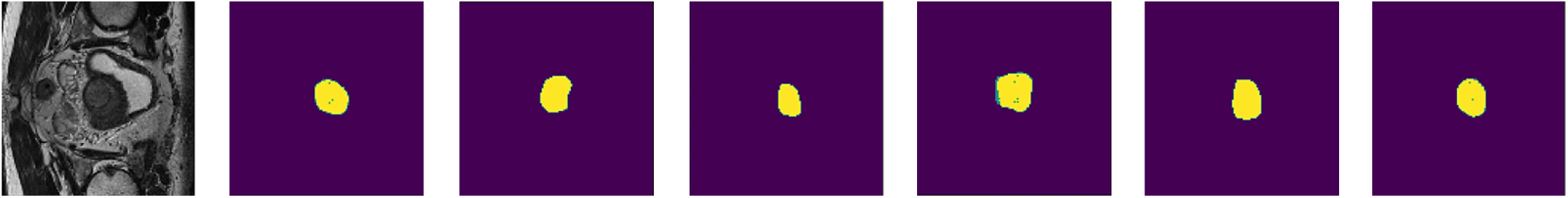
Visual Analysis.

### State of the Art Comparison

The comparison of various methods practical to prostate cancer segmentation tasks (see Table 8), highlighting the Dice coefficient and Jaccard Index (IoU) as key performance metrics. The proposed RSAUNet model, applied to 48 multi-parametric MRI studies from Radboud University, outperformed all other models with a DC of 0.998 and an IoU of 0.965. In comparison, Jiang et al.’s 2024 study using multi-scale deep supervision and TransUNet accomplished a DC of 0.939, and Zhang et al.’s 2019 bi-attention adversarial network obtained 0.859 with an IoU of 0.757. Models like SAM-UNETR from Alzate-Grisales et al. and diffusion models from Toosi et al. achieved lower DC of 0.624 and 0.532, separately, indicating the superior segmentation performance of RSAUNet. This table underscores RSAUNet’s advancement in segmentation accuracy, as evidenced by its significantly higher DC and IoU values, setting a new benchmark in prostate cancer segmentation.

**Table 8:** Comparison of RSAUNet Model with Other Techniques.

| Year/Ref | Techniques | Dataset Used | Dice Coefficient | Jaccard Index (IoU) |
| --- | --- | --- | --- | --- |
| 2024/ Jiang et al. [21] | TransUNet | Micro-ultrasound images of 75 patients | 0.939 | ----- |
| 2024/ Toosi et al. [25] | Diffusion models (DDPM), Multi-angle Maximum Intensity Projections (MA-MIP), OSEM Algorithm | 510 PSMA-PET/CT images | 0.532 | 0.433 |
| 2024/ Jeong et al. [27] | nnU-net | CT images of 18 prostate cancer patients | 0.84 | ----- |
| 2024/ Kuanar et al. [29] | nnU-Net, PSA-density (PSAD), transition zone PSAD (TZ-PSAD) | 1020 prostate MRIs for model development, 3461 additional internal test cases, and 1460 external test cases (PI-CAI challenge) | 0.91 | ----- |
| 2023/ Li et al. [30] | 3D-Mask RCNN | 133 patients (71 with prostate cancer, 62 benign tumors), T2WI MRI | 0.849 | ----- |
| 2023/ Alzate-Grisales et al. [31] | SAM-UNETR | Prostate158, PI-CAI Challenge datasets | 0.624 | 0.491 |
| 2022/ Pellicer-Valero et al. [32] | 3D Retina U-Net | ProstateX, Valencian Oncology Institute Foundation | 0.945 | ----- |
| 2022/ Duran et al. [33] | ProstAttention-Net with attention mechanism | 219 MRI (PROSTATEx-2) | 0.875 | ----- |
| 2021/ Zhang et al. [34] | SCFF | 358 MRI images | 0.657 | ----- |
| 2021/ Chahal et al. [35] | UNet with Xception in Encoder, Local Residual Connections in Decoder | PROMISE12 Dataset | 0.9750 | ----- |
| 2020/ Kiljunen et al. [36] | DL-based AST model | 30 Prostate Cancer Patients | 0.94 | ----- |
| 2019/ Zhang et al. [37] | Bi-attention adversarial network (GAN with U-Net as the backbone) | MR T2-weighted images from 120 patients from Shanghai Tenth People Hospital | 0.859 | 0.757 |
| 2018/ Hossain et al. [38] | VGG19RSeg | ----- | 0.9457 | ----- |
| **Proposed Model** | **RSAUNet** | **48 multi-parametric MRI studies provided by Radboud University, Netherlands** | **0.998** | **0.965** |

Figure 23 compares Dice Coefficients achieved by various models alongside the proposed RSAUNet model. The model outdoes the others with a Dice Coefficient of 0.998, indicating superior segmentation accuracy. Other models, like Chahal et al. (0.975) and Pellicer-Valero et al. (0.945), have also shown good performance, but not as accurate as the RSAUNet. Toosi et al. (0.532) demonstrates much lower accuracy, emphasizing how far these more recent approaches have come, especially those using residual blocks and attention mechanisms such as the RSAUNet. The comparison reflects the efficiency of the proposed model towards segmentation of prostate cancer with more accuracy.

**Figure 23:**
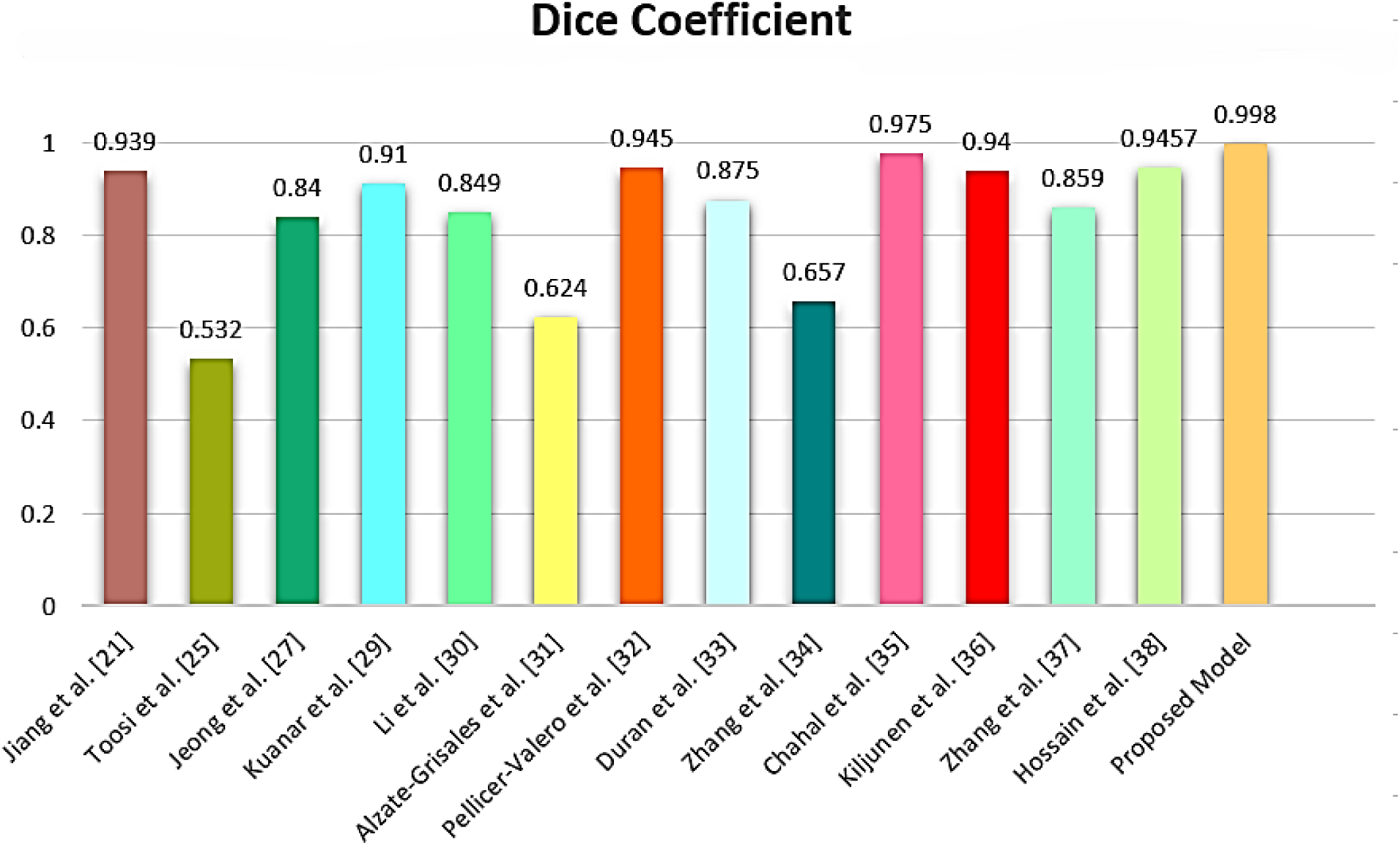
Comparison of Dice Coefficient with Proposed RSAUNet Model.

## Conclusion and Future Work

The paper presents a resilient and very accurate model called RSAUNet for prostate cancer segmentation using MRI images. The use of Residual Blocks, Swin Transformer Blocks, and Attention Mechanisms within the U-Net framework has contributed to RSAUNet’s exceptional performance in segmenting the complex anatomical features of the prostate and accurately identifying cancerous regions. The combination of Residual Blocks, Swin Transformer Blocks, and Attention Mechanisms within the U-Net architecture has demonstrated its ability to segment the intricate anatomical structure and precisely localization cancerous regions. The model was evaluated by comparison with a number of state-of-the-art segmentation approaches, consistently performing among the highest performance with Dice coefficient 0.998 and Jaccard index (IoU) of 0.965. Future studies will extend the model to incorporate the multi-modal imaging data, such as PET/CT scans, to enhance applicability to clinical settings. In addition, the applicability of RSAUNet will be explored in real time situations in the clinical environment, highlighting how computational efficiency can be improved while preserving the accuracy of the system. Overall, RSAUNet represents a significant improvement in automated medical imaging analysis. The integration into the diagnostic process could help to identify the disease earlier, treat it more precisely, and optimize the prognosis of prostate cancer.

## Declarations

### Ethical Approval

Not applicable.

### Consent to Publish declaration

The authors hereby give consent to publish the manuscript once accepted.

### Consent to Participate declaration

Not applicable

### Clinical Trial Number

Not applicable

### Conflict of Interest

The authors declare no conflicts of interest.

### Authors’ contributions

R.S., S.G., S.J. conceptualized the framework, developed the model, performed the performance analysis, and wrote the manuscript. R.S., S.G., S.J., D.G., S.Mg. also contributed to developing the model and performed the performance analysis. M.W. and S.Mk. worked on performance analysis, validation of results, manuscript review, and editing.

### Funding

The authors received no funding from their institutes for this study.

## Data Availability Statement

The dataset used in this study is publicly available at Kaggle (“Prostate annotated dataset for image segmentation”) https://www.kaggle.com/datasets/haithem1999/prostate-annotated-dataset-for-image-segmentation/data.

## Notes

### Competing Interest Statement

The authors have declared no competing interest.

